# Mapping the attractor landscape of Boolean networks

**DOI:** 10.1101/2024.09.30.615897

**Authors:** Van-Giang Trinh, Kyu Hyong Park, Samuel Pastva, Jordan C Rozum

## Abstract

Boolean networks are popular dynamical models of cellular processes in systems biology. Their attractors model phenotypes that arise from the interplay of key regulatory subcircuits. A succession diagram describes this interplay in a discrete analog of Waddington’s epigenetic attractor landscape that allows for fast identification of attractors and attractor control strategies. We present a new approach to succession diagram construction for asynchronously updated Boolean networks, implemented in the biologist’s Boolean attractor landscape mapper, `biobalm`, a Python 3 library. We compare the performance of `biobalm` to similar tools and find a substantial performance increase in succession diagram construction, attractor identification, and attractor control. To illustrate the utility of `biobalm`, we perform the most comprehensive comparative analysis to date of the succession diagram structure in experimentally-validated Boolean network models of cell processes and random ensembles. We find that random models (including critical Kauffman networks) have relatively small succession diagrams, indicating simple decision structures. In contrast, non-random models from the literature are enriched in extremely large succession diagrams, indicating an abundance of decision points in their dynamics and suggesting the presence of complex Waddington landscapes in nature.

## 1 Introduction

Biomolecular networks underpin cellular decisions and are essential in genotype to phenotype mapping. They represent the interactions between molecular entities within a cell, such as genes, proteins, and small molecules. Their kinetic parameters, however, are notoriously difficult to measure or estimate. Fortunately, living systems are often qualitatively robust to these parameters (see e.g. [40]), motivating widespread use of qualitative modeling in systems biology, with *Boolean networks* (BNs) being especially popular [18, 1, 10, 42, 35, 30]. First introduced in a gene regulatory context by Kauffman [15] as a means to study canalization (epigenetic robustness) and the emergence of phenotypic order, BNs consist of interlinked Boolean automata: each automaton’s state (ON or OFF) is dynamically updated by the states of its linked automata according to a fixed update rule. This state evolves (either *synchronously* or *asynchronously*) in discrete time steps, eventually converging to one of several *attractors* (minimal sets of states from which no escape is possible). These attractors then typically correspond to phenotypes of interest. BNs can exhibit ordered, disordered, or critical perturbation responses, which reflects the robustness of their associated biological phenotypes [6, 2, 24]. This has important basic science implications, but also biomedical significance: key *driver nodes* that disrupt undesired phenotypes represent potential drug targets, in some cases experimentally validated [36, 19, 20, 8].

One approach toward understanding phenotype robustness is through the self-sustaining configurations of small subnetworks in a BN, called *stable motifs*. These correspond to *trap spaces* within network dynamics—hypercubes in the state space from which there is no escape [43, 16]—and have analogs in ODEs [29, 28]. A *succession diagram* (SD), roughly analogous to Waddington’s canalization landscape [41], is a directed acyclic graph that describes how these trap spaces nest within one another, indicating how entering one region of the phenotypic space is predicated on (or forbidden by) entering another [43, 30]. The leaf nodes of the SD are the *minimal trap spaces*, each of which contains at least one attractor. Identifying these gives enormous computational advantages and insight into the possible behaviors of a biological system [17, 31, 39, 30].

Identifying one BN attractor (resp. all attractors) is NP-hard (resp. #P-hard) because it contains N-SAT as a sub-problem [21]. Fortunately, biologically significant BNs are often sparse, which can be leveraged in attractor identification algorithms. Still, critical bottlenecks remain that render many biologically important networks intractable. Previously, each of the authors of this work have independently explored three approaches to overcoming these bottlenecks.

First, `pystablemotifs` [31] leveraged parity and time-reversal transformations to extend and accelerate the iterative succession diagram methodology of [44] and was used to construct the first-ever exact attractor repertoires for genome-scale BNs [32]. A limitation of `pystablemotifs` is its need to frequently compute Blake canonical forms (i.e., all prime implicants) for all update rules and their negations, limiting its use to very sparse networks where such computations are easy. Following `pystablemotifs`, `AEON.py` [3] was released, using binary decision diagrams along with transition guided reduction [4] to dramatically improve the efficiency of graph exploration in attractor identification. Finally, and most recently, `mts-nfvs` was released [39]. It uses an alternate scheme, implemented within the `trappist` library [38], to identify trap spaces via Petri-net encodings, which are easier to compute than the Blake canonical form. Furthermore, it leverages properties of negative feedback vertex sets (NFVS) to more efficiently search for motif-avoidant attractors [12, 11]. It uses minimal trap spaces and preprocessing heuristics to simplify or avoid reachability analysis in most cases. Still, non-minimal trap spaces and their nesting relationships further improve the method.

Each of these three methods is faster than the last. The algorithm we present here incorporates advantages from each, along with new insights about how to efficiently build SDs, resulting in `biobalm`. It uses the iterative SD approach of `pystablemotifs`, efficient rule representation and symbolic state-space searching from `AEON.py`, and the trap space identification method and NFVS approach of `mts-nfvs`. We demonstrate substantial speed improvement compared to these prior methods. Our method enables systematic exploration of motif-avoidant attractors in large BNs, exact attractor identification and control in previously intractable experimentally-supported BNs, and analysis of SD scaling in random and non-random BNs to provide insight into the emergence of canalization in biology.

## 2 Background

Here, we give an overview of key Boolean modeling concepts and notation. More formal details are given in Supplementary Text S1.

### 2.1 Boolean networks

An *asynchronous Boolean network* (ABN) of dimension *n*, denoted *B*, is a non-deterministic dynamical system. *States* of *B* are *n*-dimensional Boolean vectors *x* ∈ 𝔹^*n*^, with *x*_*v*_ denoting individual vector components. Each network variable *v* is assigned a Boolean *update function f*_*v*_ : 𝔹^*n*^ → 𝔹 that governs its time evolution. At each discrete time step, the value of a non-deterministically selected variable is updated to match the output of its update function. When the variables are indexed, we may write *f*_*i*_ to refer to 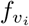 and *x*_*i*_ to refer to 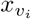. An example ABN with *n* = 4 is shown in Figure 1a.

**Figure 1:**
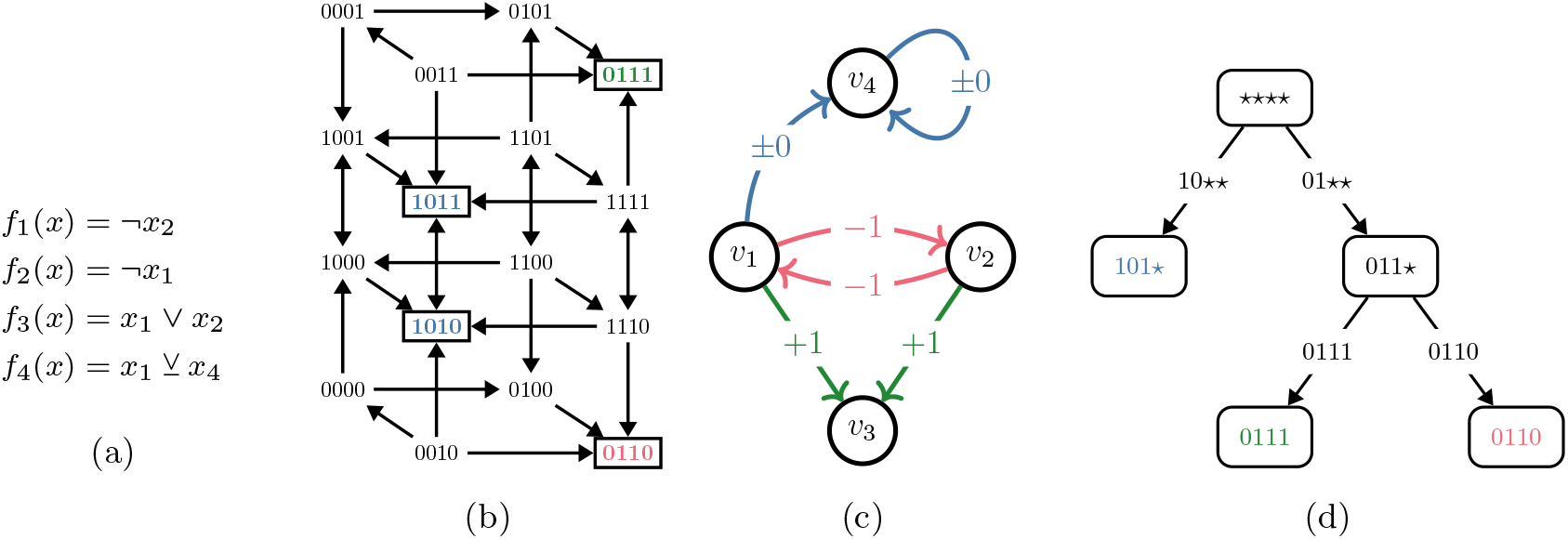
Illustration of the core concepts utilized in this paper. (a) A simple ABN *B* = {*f*_1_, *f*_2_, *f*_3_, *f*_4_}, with ¬ denoting negation, ∧ conjunction, ∨ disjunction and ⊻ the exclusive or. (b) *STG*(*B*). Self-loops are omitted for visual clarity. Attractor states are highlighted and distinguished by color. (c) *IG*(*B*). Signs +1 (green), − 1 (red), and *±* 0 (blue) denote positive, negative, and non-monotonic influence, respectively. (d) *SD*(*B*). Edges are labeled with the maximal trap spaces that percolate to the target nodes. As is typical (but not guaranteed), the minimal trap spaces (leaf nodes) and attractors (in (b)) coincide.

The dynamics of ABN *B* are encoded in a *state transition graph STG*(*B*) whose nodes are the states *x* ∈ 𝔹^*n*^ of *B*. An edge *x* → *y* exists in *STG*(*B*) if and only if *B* can update from state *x* to state *y* in one time-step (i.e., *f*_*v*_(*x*) = *y*_*v*_ for some *v*). The *STG*(*B*) corresponding to the network from Figure 1a is shown in Figure 1b. The core feature of each *STG*(*B*) are its *attractors*: minimal subsets of 𝔹^*n*^ that are closed under time evolution. These are also highlighted in Figure 1b. Note that other BN *update schemes* exist, such as the synchronous update (see [24] for a detailed discussion). However, much of the contributions of this paper do not depend on the chosen update scheme (further discussion is given in Supplementary Text S1.2).

ABN dynamics can be viewed as arising from a network of interactions among Boolean automata called an *influence graph* (IG), denoted *IG*(*B*), with nodes *v*_1_, …, *v*_*n*_. An edge from *v*_*i*_ to *v*_*j*_ indicates that the state *x*_*i*_ of automaton *v*_*i*_ is a non-redundant input^1^ to the update function *f*_*j*_. The sign of an edge from *v*_*i*_ to *v*_*j*_ can be −1 for inhibition, +1 for activation, or *±* 0 if the impact of *v*_*i*_ depends on the remaining regulators. Figure 1c shows *IG*(*B*) for the example network from Figure 1a. The IG is very useful for model analysis, as complex dynamics arise from the interplay between positive and negative feedback loops in the IG.^2^ It is therefore often advantageous to identify a *feedback vertex set* (FVS), which is a set of nodes that intersects every cycle of *IG*(*B*). Controlling the activation of an FVS is sufficient to drive an ABN into any of its attractors [7, 45]. Furthermore, a *negative* FVS (NFVS) is a set of nodes that intersects every negative feedback loop. Fixing the nodes of any NFVS ensures that all variables in an ABN eventually stabilize [12]. Typically, we are interested in an (N)FVS with as few nodes as possible. Identifying the true minimum (N)FVS is computationally difficult, but in our method, a heuristic estimate is sufficient (computed using [3]). The IG in Figure 1c has two minimum FVSs: {*v*_1_, *v*_4_} and {*v*_2_, *v*_4_}. Meanwhile, the only minimum NFVS is {*v*_4_}.

### 2.2 Network trap spaces and succession diagrams

Network *subspaces* represent special subsets of 𝔹^*n*^ given by fixing some of the variables. Formally, subspaces are the members of 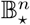, where 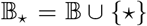. The value of a network variable *v* in a subspace 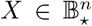 can be either fixed (*X*_*v*_ = 0 or *X*_*v*_ = 1), or free (*X*_*v*_ = ⋆). Each subspace *X* corresponds to a set of states 𝒮 (*X*) ⊆ 𝔹^*n*^ that agree with *X* in all fixed variables. For example, *X* = 011⋆ corresponds to 𝒮 (*X*) = 0110, 0111. We refer to *X* and 𝒮 (*X*) interchangeably as context allows.

*Trap spaces* are subspaces that are closed under time evolution. Of special importance are (inclusion) *minimal trap spaces*, as each is guaranteed to contain at least one attractor. Importantly, however, attractors can also appear outside of the minimal trap spaces. For example, consider a simple ABN *f*_1_(*x*) = *f*_2_(*x*) = *x*_1_ ⊻ *x*_2_. This network has two trap spaces, ⋆⋆ and 00, of which 00 is minimal. However, the network also has two attractors: *A*_1_ = {00} and *A*_2_ = {01, 10, 11}, meaning that *A*_2_ does not lie within any minimal trap space. Such attractors are called *motif-avoidant* [27]. When a motif-avoidant attractor exists, the set of minimal trap spaces is called *incomplete* [16, 17]. *Motif-avoidant attractors, as noted by [32, 24] and in Section 4.1, are rare, but it is very difficult to rule out their existence a priori*. Various methods have been devised for detecting whether a set of minimal trap spaces is incomplete [17, 32, 11].

The subspace *percolation* is the process of propagating fixed values among the network variables. We write 𝒫 (*X*) to denote the one-step percolation of the subspace *X*. This updates each variable *v* which is free in *X* (i.e. *X*_*v*_ = ⋆) to a fixed value *b ∈* 𝔹 if and only if *f*_*v*_(*x*) = *b* for every state *x* in *X*. Repeatedly applying the 𝒫 operator (up to *n* times) results in a subspace where no further variables can be updated, denoted 𝒫^∞^(*X*). We say that *X percolates* to the subspace 𝒫^∞^(*X*) and that *X* is *percolated* if *X* = 𝒫^∞^(*X*). For example, consider the ABN from Figure 1a and the subspace 𝒫 *X* = ⋆0⋆⋆. This one-step percolates to 𝒫 (*X*) = 10⋆⋆ because *f*_1_(*x*) = *¬x*_2_, which is equivalent to *¬*0 for every network state in *X* (since *x*_2_ = 0 for every *x ∈ X*). Meanwhile, for *f*_3_ and *f*_4_, both output values are possible in *X*, hence they remain free. A second application of 𝒫 yields the subspace 101⋆ because *f*_3_(*x*) = *x*_1_ ∨ *x*_2_ is equivalent to 1∨0 for every state in 10⋆⋆. Further applications of 𝒫 result in no additional changes, because *f*_4_(*x*) simplifies to *¬x*_4_ in 101⋆. Therefore 𝒫^∞^(*X*) = 𝒫^2^(*X*), i.e. ⋆0⋆⋆ percolates to 101⋆.

Knowledge of network’s trap spaces and their relationships with one another can aid in understanding the network’s long-term dynamics, including its attractors [39, 31, 30] and response to interventions [43, 44, 32]. Percolated trap spaces are typically emphasized because any state in *X* eventually evolves to a state in the percolation of *X*. To formalize relationships between percolated trap spaces, [43] introduced *succession diagrams*. A succession diagram^3^ of an ABN *B*, denoted *SD*(*B*), is a rooted, directed acyclic graph. The vertices of *SD*(*B*) are exactly all percolated trap spaces of *B*, with the edge relation describing how these nest within one another (by set inclusion). The root node is the percolation of ⋆^*n*^.

Notice that the terminal (leaf) nodes of *SD*(*B*) are exactly the minimal trap spaces of *B*. Furthermore, the successors of a node *X* correspond to the trap spaces obtained by percolating trap spaces that are subset-maximal within *X*. Borrowing terminology from related hypergraph structures [43, 32], we call such maximal trap spaces *stable motifs* and show them as edge labels when presenting succession diagrams. Most often, every maximal trap space percolates to a distinct succession diagram node. However, in some cases, multiple maximal trap spaces percolate to the same subspace, in which case the edge can be annotated with multiple stable motifs. The succession diagram of the network from Figure 1a is depicted in Figure 1d.

#### Control interventions

Succession diagrams are also useful for attractor control, and form the basis for several ABN control algorithms [44, 31]. The majority of these methods involve selecting a path in *SD*(*B*) from the root node to a target trap space containing the desired attractor. At each branch point along the path, an intervention is selected to ensure that the system will eventually enter the trap space corresponding to the selected path. The union of these interventions drives the system to the target trap space with probability 1. The advantage of this approach is that it subdivides the attractor control problem into smaller, more manageable pieces. Typically, controlling entry into each trap space along the selected path involves fixing only a small subset of the Boolean variables. In `biobalm`, we have implemented two control algorithms from [31] with only slight modifications to allow for dynamic expansion of the succession diagram (see Supplementary Text S4 for details).

## 3 Algorithm

In this section we give an overview of the methods implemented by `biobalm`. Details, pseudo-code, and proofs are given in Supplementary Text S2-5.

### 3.1 Succession diagram construction

We introduce several innovations to the algorithms of [43] and [32, 31]. At a high level, `biobalm` is broadly similar to the previous tools (Supplementary Text S2.3): the root node 𝒫^∞^(⋆^*n*^) of the ABN *B* is established by percolating the trivial trap space ⋆^*n*^, and it is stored in a digraph *SD*(*B*). Then, *SD*(*B*) is further expanded by selecting a node (percolated trap space) *X* in *SD*(*B*), identifying all maximal trap spaces within *X*, and percolating them to obtain the successor nodes. These are incorporated into *SD*(*B*). Once the maximal trap spaces are computed for all nodes, the digraph *SD*(*B*) is the succession diagram of *B*, as introduced in Section 2.

Compared to previous tools, however, `biobalm` has several key advantages. First, we use a more efficient trap space identification method [11, 39], which we further improved by implementing a heuristic for achieving more compact encodings of update functions (Supplementary Text S2.1). Second, we have implemented a more efficient percolation function that avoids the expensive step of recomputing prime implicants (Supplementary Text S2.2). Third, we adapt the attractor identification method of [39] to apply to arbitrary percolated trap spaces instead of only minimal trap spaces (Supplementary Text S3.3). Finally, we decouple attractor identification from SD construction and implement schemes for partially expanding the SD (discussed further in Section 3.2).

#### 3.1.1 Partial expansion strategy

Previous methods [43, 31] construct *SD*(*B*) by preferentially expanding deeper nodes. In `biobalm`, we implement multiple strategies, including depth-first and breadth-first expansion, allowing early termination if certain size or depth threshold is exceeded. We also developed several partial expansion strategies, which produce a sub-graph of *SD*(*B*) in which certain nodes are not expanded, meaning their child nodes in *SD*(*B*) are omitted. Upon completion, these methods produce partial SDs that are functionally equivalent to the full SD for attractor identification and control, but which eliminate computationally expensive (and cognitively burdensome) redundancies.

Of these strategies, the one that removes the most redundancies, and which we have selected as the default method for `biobalm`, is the *source block* expansion (Supplementary Text S5). Conceptually, this method is similar to [37, 14] in that it identifies hierarchies of sub-networks (*blocks of variables*) that are mutually independent and can be processed separately, thereby eliminating redundancy associated with permuting the order of trap space entry. Full details are given in Supplementary Text S5.

In Figure 2, we show the application of this method to the mammal sex-determination network developed in [34] with input values *true* (see also Supplementary Text S8.2). In the root node of the SD, there are six maximal trap spaces divided among four source blocks: *C*_1_ = {*v*_8_, *v*_9_}, *C*_2_ = {*v*_1_, *v*_2_}, *C*_3_ = {*v*_7_, *v*_8_, *v*_9_}, and *C*_4_ = {*v*_1_, …, *v*_9_}. Initially, we choose *C*_1_ (blue), which in one branch also eliminates *C*_3_. In the other branch, we then pick *C*_3_ (green), which is now reduced to just *v*_7_ (*v*_8_ and *v*_9_ are already fixed by *C*_1_). Next, we expand *C*_2_ (red). This either leads directly into a minimal trap space, or it simplifies *C*_4_ to just two (purple) or three (yellow) variables, depending on the choice in *C*_1_. Notice that we expanded much fewer SD nodes (17) compared to the full SD (36). Also, the largest network ever considered in the attractor identification has only three variables instead of nine.

**Figure 2:**
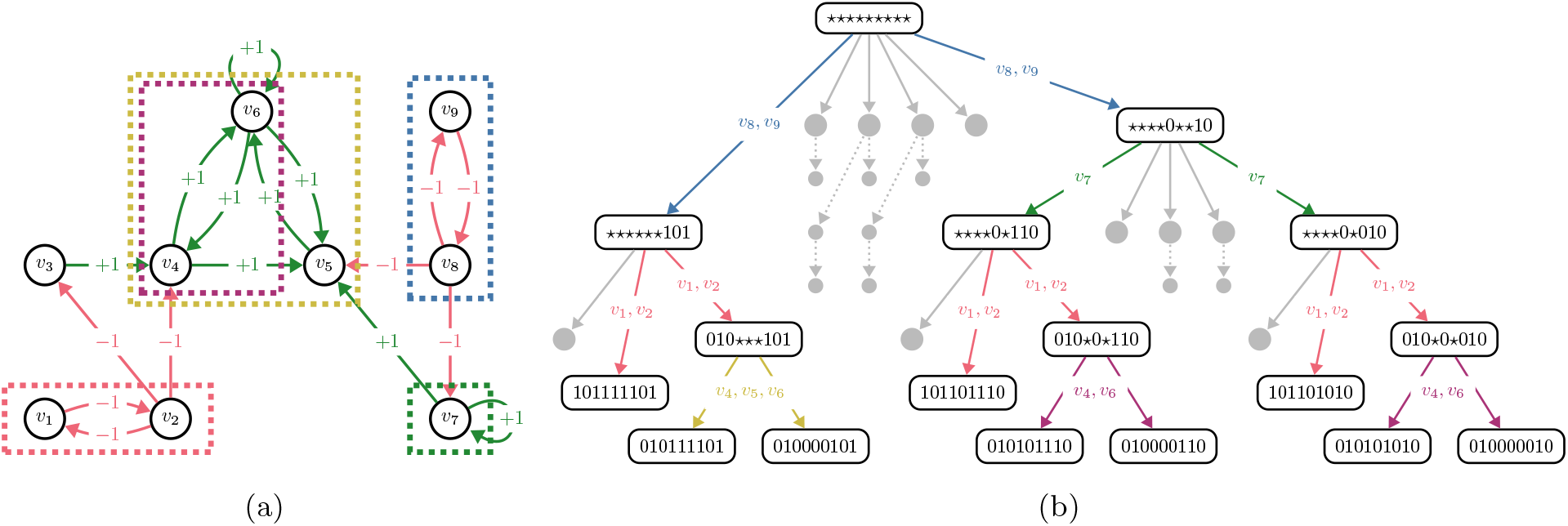
Example of a partial SD expansion by independent source blocks: (a) The IG of the simplified BN from [34] (see Supplementary Text S8.2 for details), with relevant variable blocks highlighted; (b) The partially expanded succession diagram. Colored edges are labeled with the expanded source blocks. Larger grey nodes represent trap spaces that are discovered but never expanded. Smaller grey nodes are never discovered, but appear in the full SD. For brevity, only one edge is shown per each non-expanded node.

### 3.2 Attractor identification

In `biobalm`, we consider two variants of the attractor identification problem: First, the *attractor sets* problem is to determine every set of states that represents an attractor of network *B*. This is how the attractor identification is typically understood, but it means that each attractor set must be sufficiently small such that it is fully identifiable. As this is not always the case for large complex attractors, we also consider the *attractor seed* problem, which is to identify exactly one representative *seed* state for each attractor. This allows us, for example, to identify the presence of motif-avoidant attractors without fully enumerating their states.

Our approach in `biobalm` is based on the method of `mts-nfvs` [39], which we have significantly improved and extended to apply to arbitrary percolated trap spaces rather than only the minimal ones. Our workflow applies to each SD node *X* individually. A summary is given in Figure 3 as well as Supplementary Text S3.

**Figure 3:**
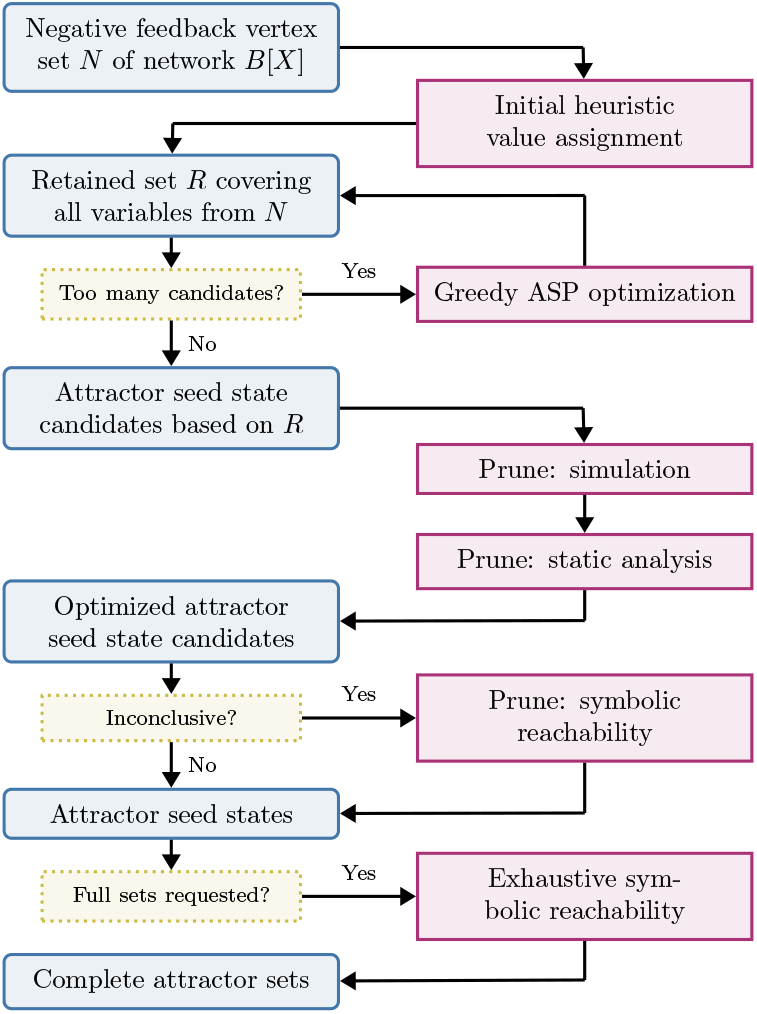
Attractor identification workflow for a fixed node (trap space) *X* of the succession diagram *SD*(*B*). Blue and purple boxes represent data and computational steps, respectively.

Recall that attractors cannot cross trap space boundaries, and that the 𝒯 (*A*) (smallest trap space that contains *A*) for each attractor *A* is always a fully percolated trap space. Thus it is sufficient to search for attractors only within percolated trap spaces, i.e., the nodes of *SD*(*B*). Furthermore, when searching an SD node *X* for attractors, we can disregard not only the states outside *X*, but also the states in each successor node *Y* (as these are considered separately when searching *Y*). We use the NFVS-based method of [39] to identify candidate attractor seeds in each percolated trap space, and then verify or eliminate them using randomized simulation, static analysis (using `pint` [26]), or symbolic reachability (using `AEON.py` [3]).

## 4 Results

### 4.1 Benchmarks

To evaluate the overall effectiveness of `biobalm`, we consider the attractor seed identification problem over a large collection of real-world (230 networks with 14,010 input configurations; from the BBM dataset [25]) and synthetic (2760 networks; critical N-K, nested canalyzing, and dense ensembles) Boolean networks. A detailed description of experiment setup is given in the Supplementary Text S6. We compare `biobalm` with `AEON.py` [3] and `mts-nfvs` [39]. Figure 4 summarizes the attractor benchmark results. The top panel shows the number of benchmarks completed as a function of time. The bottom panel compares the performance on individual benchmarks. In Supplementary Text S6, we also provide an extended versions of Figure 4, stratified across individual tools and network ensembles. Importantly, we have not encountered any motif-avoidant attractor in either of ensembles.

**Figure 4:**
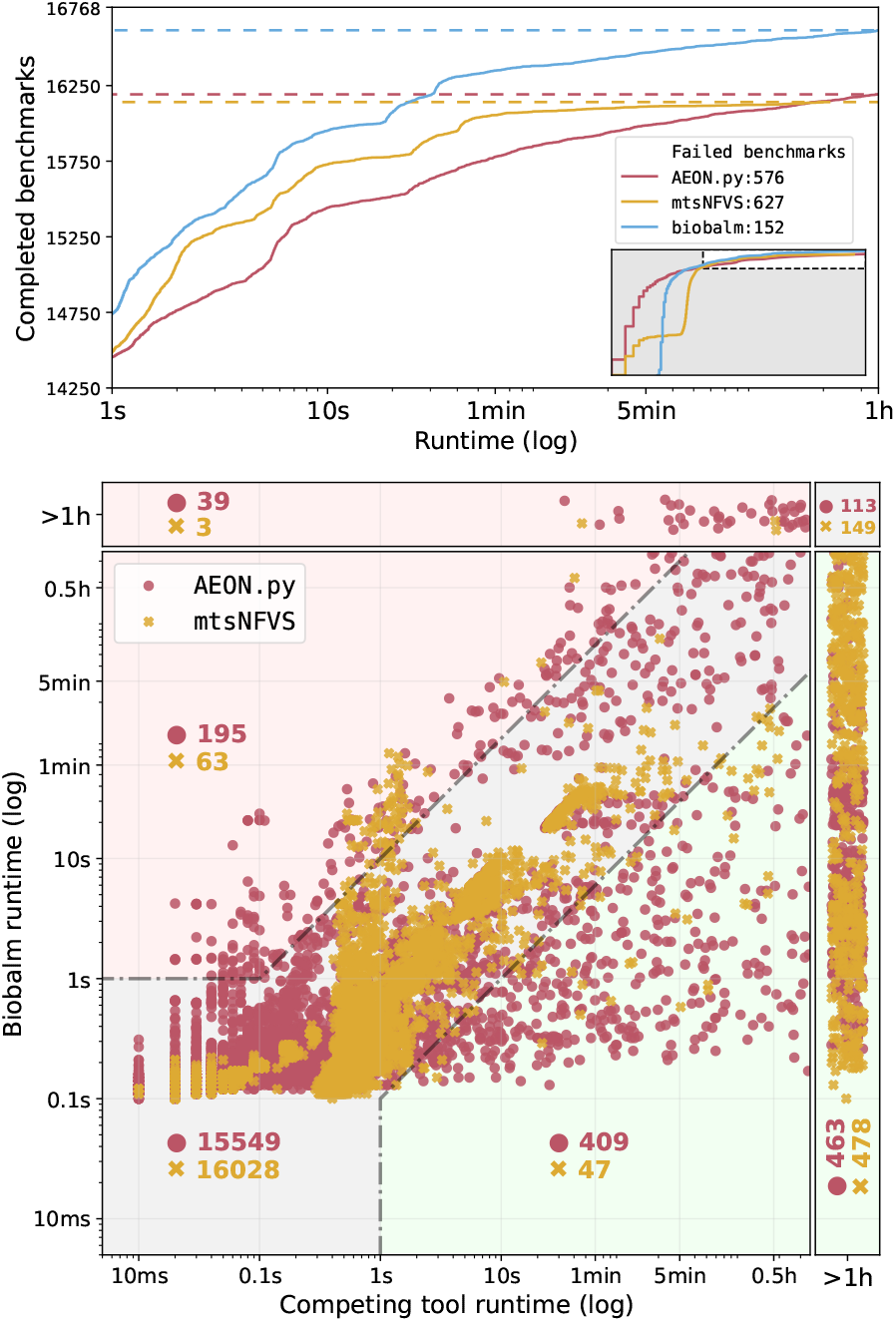
Performance comparison of `biobalm` versus `AEON.py` and `mts-nfvs` for attractor identification. (Top) The total number of completed benchmarks within `1h` timeout (vertical axis) out of our test ensemble of 16,768 benchmark models as a function of time (horizontal axis, logarithmic scale). We show times `>1s` in the main panel; the full range is shown in the inset and in Supplementary Text S6. (Bottom) Runtime of individual benchmark instances. Timeouts are placed at the margins of the plot, indicated by the “`>1h`” labels. Green (red) regions represent results where `biobalm` was at least 10*×* faster (slower) than the competing tool and the slower tool took longer than `1s` to complete. The grey region contains instances where tools performed similarly. The number of cases that fall into each region is indicated in red or yellow text for `AEON.py` and `mts-nfvs`, respectively.

We have not included `pystablemotifs` in the attractor identification benchmarks because [3] demonstrated that `AEON.py` is superior for this task. However, we have tested `pystablemotifs` against `biobalm` on a smaller set of real-world models to evaluate succession diagram expansion and control. Results are provided in Supplementary Figure S9 and S10. In these tests, `biobalm` completed almost all benchmarks at least 10*×* faster than `pystablemotifs`.

### 4.2 Attractor landscape ensembles

To demonstrate the utility of `biobalm`, we have compared the full succession diagram structure of 230 ABN models of cell processes from the BBM dataset [25], in 14,010 parameter configurations, to the succession diagrams of 69,000 random ABNs drawn from three null model ensembles. The null model ensembles include two ensembles of critical N-K models [15] with in-degree K=2 and K=3, and an ensemble of nested canalizing function (NCF) networks generated using the methods of [22]. The NCF networks have nested canalizing regulatory functions—meaning inputs determine outputs in a hierarchical manner—and topology matched to biological networks as reported by [13]. We generated 100 random networks with equal size matched to each BBM model, thereby resulting in three ensembles of 23,000 networks each.

As shown in Figure 5, the distribution of succession diagram sizes for empirical models is more heterogeneous and has a longer tail than for our three ensembles of random networks. The mean, variance, and kurtosis of the BBM distribution are statistically significantly higher than for random networks (via bootstrapped 95% confidence intervals–see Supplementary Text S7.3). We observe a similar pattern in the number of attractors (see Supplementary Table S2). Additionally, we controlled for any automatically-generated models within BBM and observed similar results regardless of the model origin (Supplementary Text S7.4).

**Figure 5:**
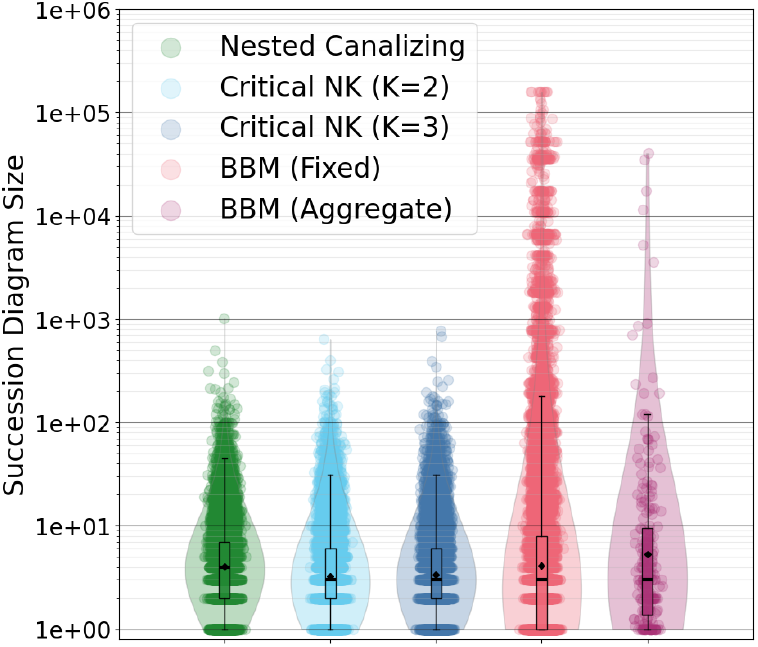
Distributions of succession diagram size (number of nodes in the fully-expanded succession diagram) for various ABN ensembles. BBM networks with random input configurations (up to 128 independent samples) are shown in red, with the average succession diagram size across these samples for each network depicted in purple. Random networks of different types are indicated in green, cyan, and dark blue, and are constructed to match the distribution of the number of variables in the BBM ensemble. Gaussian noise is added to the horizontal position of each point. Box plots and density plots are computed in log space.

We observe that succession diagram size and the number of attractors grow exponentially with succession diagram depth (denoted *d*) in all four ensembles (Supplementary Figure S13). Note that the succession diagram is free to have any depth, any size, and any number of attractors. Succession diagram size is roughly bounded by 3^*d*^, and the number of attractors is roughly bounded by 2^*d*^, scaling bounds that correspond to networks of independent bistable cycles. Moreover, SD size as a function of depth is approximately bounded between the scaling obtained for uniform trees with three children and one child (linear chains), with many SDs falling near the scaling law obtained for uniform binary trees (two decisions for each SD node). These observations suggest that attractor landscapes are typically not broad and shallow–instead, attractor commitment arises from a series of decisions between a small number of possible choices.

## 5 Discussions

Extracting biological insights from a Boolean network requires a strong understanding of its range of possible behaviors, as well as of the circumstances under which they arise. Despite recent major advances, the problem of identifying all the attractors in a large, dense ABN remains difficult. This problem is increasingly important as modelers of cell processes develop ever larger and more complete networks (and integrate them into population-level models). We have previously contributed to this problem individually [31, 3, 39], and here, we present a combined and significantly improved approach in the Python 3 library `biobalm`, the biologist’s Boolean attractor landscape mapper.

The attractor identification benchmarks we have presented demonstrate a substantial improvement over `AEON.py` [3] (which has been previously demonstrated to significantly outperform `pystablemotifs` [31]) and a moderate speed improvement over `mts-nfvs` [39]. Crucially, `biobalm` successfully analyzed 463 networks and 478 networks where `AEON.py` and `mts-nfvs` failed, respectively. Out of these, 75 networks were uniquely solved by `biobalm` (27 biological and 48 random). With a more generous timeout of two days, `biobalm` only fails 26 benchmarks, primarily in large models with more than 100,000 complex attractors that are limited by the time needed to enumerate individual attractors. Overall, `biobalm` is overall the fastest and most robust among the tested tools. Moreover, `biobalm` is simultaneously computing the succession diagram, and therefore yields a much more informative output that describes the decision points in the network circuitry and can be used to perform computationally efficient attractor control using the control algorithms introduced in [31]. Compared to `pystablemotifs`, this results in a significant (10*×* or better) speed-up in control strategy identification.

Importantly, `biobalm` provides a modular approach to both succession diagram construction and attractor identification, enabling multiple methods and heuristics informed by the succession diagram (Figure 3). This allows to easily replace components (NFVS computation, minimal trap space computation, symbolic reachability, etc.) or introduce new methods as improvements become available.

In addition to benchmarks, we have used `biobalm` to study the distribution of succession diagram sizes in biological models from the Biodivine Boolean Models (BBM) repository [25], the largest curated collection of biological Boolean networks currently available. We compared these models to similarly sized critical Kauffman networks [15] (K=2 and K=3) as well as to an ensemble of canalizing random networks generated using the method of [23]. We observe that succession diagram depth scaling is consistent with attractor landscapes constructed from a series of small decisions, lending support to the hypothesis that biomolecular networks exhibit modular, hierarchal structure [33, 24]. We also find that the distribution of succession diagram size for published Boolean network models with fixed inputs is highly heterogeneous and contains extremely large succession diagrams (several thousand nodes), as compared to the succession diagram distributions for three biologically-inspired random model ensembles. Allowing inputs to vary in the published models would even further accentuate these differences. This suggests that, even accounting for network size, degree distribution, and the prevalence of canalizing functions in biological Boolean networks [13], they exhibit more complicated Waddington canalization landscapes at the system-level. Because the trap spaces that compose the succession diagram arise from positive feedback loops [43], we propose that non-local topological features of the influence graph (such as prevalence and overlap of cycles) play a significant role in the emergence of complex Waddington landscapes. It seems likely that local features (such as various measures of canalization in individual update functions [5]) are not sufficient to explain or predict the complexity of the attractor landscape—though they may still be informative in computing non-local metrics, e.g., via the effective graph [9]. Testing our hypothesis requires a deeper analysis of this phenomenon and remains as future work.

## Ackowledgements

We thank Réka Albert for her helpful advice regarding the development of a new SD expansion strategy. We thank Luis M. Rocha for insightful discussions regarding the role of canalization in the results presented here.

## Data and software availability

The tool `biobalm` is available online at https://github.com/jcrozum/biobalm. Further data, scripts for testing, analysis and figure generation are available online at https://github.com/jcrozum/biobalm-analysis and in the reproducibility artefact at https://doi.org/10.5281/zenodo.13854760. The Supplementary Text is available online through *Bioinformatics*.

## Footnotes

1 In some applications, redundant edges added to *IG*(*B*) may be informative, such as those that encode well-known interactions that are non-functional within the modeling context. Here however, we assume that *IG*(*B*) only consists of essential interactions and is fully determined by *B*.

2 Feedback loop sign is determined by the product of its edge signs. Cycles of sign 0 are treated as both positive and negative, because they may act as either, depending on where in the state space they are evaluated.

3 We use a novel, simplified definition of succession diagrams. Supplementary Text S1.3.3 gives a detailed comparison to the previous material on this topic.

