## Supplementary Text for "Mapping the attractor landscape of Boolean networks"

Van-Giang Trinh

Kyu Hyong Park

Samuel Pastva

Jordan C Rozum

December 19, 2024

This supplementary material serves to provide additional details that clarify the contents of the main paper. In particular, it provides exact formal definitions, algorithms and proofs for the methodology that is presented in the paper. Furthermore, it outlines the technical details of the performance evaluation and data analysis. The document is structured as follows:

- Section S1 defines commonly known concepts, such as Boolean networks, influence graphs, trap spaces, attractors, succession diagrams, and the Petri net encoding of Boolean networks.
- Section S2 formalizes the methods necessary for efficient succession diagram construction, such as percolation and trap space detection.
- Section S3 details the novel attractor detection method that follows the structure of a fully expanded succession diagram.
- Section S5 details the partial expansion of a succession diagram based on independent maximal trap spaces and proves that it is sufficient to fully identify network attractors.
- Section S6 describes the benchmark conditions and provides detailed performance results.
- Section S7 gives additional details on the analysis performed on the properties of succession diagram ensembles.

For relevant data and code, please consult the repository of the tool `biobalm` at `https://github.com/jcrozum/biobalm`, and the reproducibility artefact at `https://doi.org/10.5281/zenodo.13854760`.

### S1 Formal Preliminaries

#### S1.1 Notation

In the following, we consider *true* and *false* to be respectively interchangeable with 1 and 0 and write $\mathbb{B}$ to denote the Boolean domain $\mathbb{B} = \{0, 1\} \equiv \{\mathit{false}, \mathit{true}\}$. We also define an *extended* Boolean domain $\mathbb{B}_\star = \{0, 1, \star\}$, where $\star$ is a stand in for an *unknown* Boolean value. Finally, we define $\mathit{Var}_n = \{v_1, \ldots, v_n\}$ to be an indexed set of $n$ variables. In general, these are interchangeable with the integer indices $[1, n]$, but can be used in places where an index would be confusing (e.g., as propositions in logical formulae).

##### S1.1.1 Boolean vectors and spaces

Given a fixed $n \in \mathbb{N}$, $x \in \mathbb{B}^n$ is a Boolean *vector*, with $x \equiv (x_1, \ldots, x_n) \equiv (x_{v_1}, \ldots, x_{v_n})$ where $v_i \in \mathit{Var}_n$. For convenience, we write concrete vectors simply as strings of values (e.g., 0110 instead of $(0, 1, 1, 0)$).

In the context of Boolean networks, we typically refer to the members of $\mathbb{B}^n$ as Boolean *states*, with $\mathbb{B}^n$ being the *state space*. Furthermore, we say that any vector $X \in \mathbb{B}^n_\star$ corresponds to a *subspace* of $\mathbb{B}^n$, written $\mathcal{S}(X)$ and defined as

$$\mathcal{S}(X) = \{x \in \mathbb{B}^n \mid x_v = X_v \iff X_v \neq \star\}. \tag{1}$$

Because the Boolean subspaces of $\mathbb{B}^n$ are in exact correspondence with the vectors of $\mathbb{B}^n_\star$, we apply an abuse of notation in which $X$ can refer to $\mathcal{S}(X)$ when the context allows it. For example, we can write $0110 \in 0{\star}1{\star}$, meaning that the state 0110 belongs to the subspace $\mathcal{S}(0{\star}1{\star}) = \{0010, 0011, 0110, 0111\}$. When referring to a subspace $X$, we can also write $\mathit{fixed}(X)$ and $\mathit{free}(X)$ to denote the subset of $\mathit{Var}_n$ where $X_v \in \mathbb{B}$ and $X_v = \star$, respectively. That is, for $X = 0{\star}1{\star}$, we have $\mathit{fixed}(X) = \{v_1, v_3\}$ and $\mathit{free}(X) = \{v_2, v_4\}$.

We also define the reassignment operation $[v \leftarrow b]$ acting on $X \in \mathbb{B}^n_\star$ by

$$(X[v \leftarrow b])_u = \begin{cases} b & u = v \\ X_u & \text{else} \end{cases}, \tag{2}$$

with an analogous operation defined in the same way on $x \in \mathbb{B}^n$. Intuitively, $X[v \leftarrow b]$ represents a copy of $X$ in which the value of variable $v$ is set to the value $b$ and all other variables remain unchanged.

Finally, we apply the same notation to denote variable substitution in Boolean functions. Given $f : \mathbb{B}^n \to \mathbb{B}$, $f[v_i \leftarrow b]$ denotes a function where the input $v_i \in \mathit{Var}_n$ is substituted for a constant $b \in \mathbb{B}$:

$$f[v_i \leftarrow b](x_1, \ldots, x_n) = f(x_1, \ldots, x_{i-1}, b, x_{i+1}, \ldots x_n) \tag{3}$$

Note that the domains of both $f$ and $f[v_i \leftarrow b]$ are the same (i.e. $\mathbb{B}^n$), but the output of $f[v_i \leftarrow b]$ does not depend on the input $v_i$.

#### S1.2 Boolean networks

A Boolean network associates $n$ Boolean variables to $n$ Boolean update functions:

**Definition S1.1** *A* Boolean Network (BN) *over $Var_n$ is a collection of* update functions $B = \{f_1, \dots, f_n\}$*, such that each $v \in Var_n$ is associated with exactly one update function $f_v : \mathbb{B}^n \to \mathbb{B}$.*

**Example S1.1** *Consider a Boolean network $B = \{f_1, f_2, f_3, f_4\}$ over $Var_4$ s.t.:*

$$
\begin{aligned}
f_1(x) &= \neg x_2 \\
f_2(x) &= \neg x_1 \\
f_3(x) &= x_1 \vee x_2 \\
f_4(x) &= x_1 \veebar x_4
\end{aligned}
$$

*We will refer to this BN as a running example in the following text. In this notation, $\neg$, $\wedge$, and $\vee$ represent Boolean negation ("NOT"), conjunction ("AND"), and disjunction ("OR"), respectively. The exclusive-or operator $\veebar$ is defined such that $x_1 \veebar x_4 \equiv (x_1 \wedge \neg x_4) \vee (\neg x_1 \wedge x_4)$.*

The update functions, together with a *variable update scheme*, describe the discrete time evolution of a state vector. Here, the update scheme chooses at each time-step $t$ a subset of variables $S^t \subseteq Var_n$ that is to be updated.

The two most commonly used update schemes are the *synchronous* and the general *asynchronous* update schemes. The simplest case is the synchronous update scheme, in which all nodes are updated to match the output values of their update functions at every time step (i.e. $S^t = Var_n$ for all $t$), resulting in a deterministic dynamical system. Synchronous update is widely used because of its mathematical and computational simplicity, but it also has a tendency to give rise to spurious oscillations and it necessarily treats all update events as occurring on the same time scale. Meanwhile, the *general asynchronous* update scheme, was introduced to address these difficulties (at the cost of determinism), and gives rise to an *Asynchronous Boolean Network (ABN)*:

**Definition S1.2** *An* Asynchronous Boolean Network (ABN) *is a BN over $Var_n$ in which a Boolean vector $x^t$ at time $t$ is updated to a Boolean vector $x^{t+1}$ at time $t+1$ by non-deterministically selecting a single variable $v$, and applying $x^{t+1} = x^t[v \leftarrow f_v(x^t)]$. The non-deterministic selection process is constrained only in that the selection of any variable must be possible.*

Here, a single update function is applied in every step to produce the new state (i.e., $|S^t| = 1$). As such, the behaviour of the network is non-deterministic based on the choice of the applied update function. This has the effect of disrupting oscillations that arise from positive feedback loops. By assigning a numeric probability of selecting each variable for update, time scale differences can be incorporated [1].

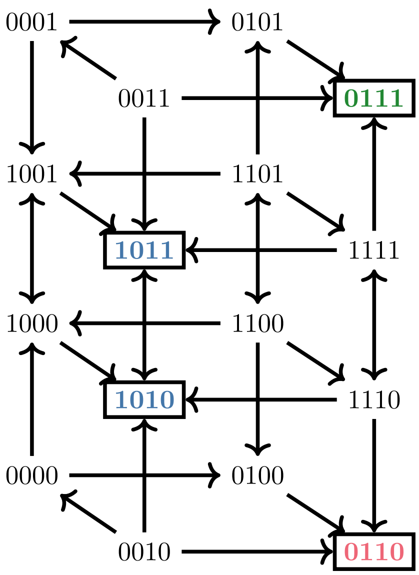


Figure S1: Asynchronous STG of the BN from Example S1.1 (rules $f_1(x) = \neg x_2$; $f_2(x) = \neg x_1$; $f_3(x) = x_1 \vee x_2$; $f_4(x) = x_1 \veebar x_4$). Vertices are network states (members of $\mathbb{B}^4$). Self-loops are omitted for visual clarity. Attractor states are highlighted with a box, and the three attractors are distinguished by text color.

All possible trajectories of an ABN can be described by its State Transition Graph (STG), whose vertices are Boolean vectors and whose edges consist of all pairs $(x, y)$ such that setting $x^t = x$ and updating some variable $v$ results in $x^{t+1} = y$; formally:

**Definition S1.3** *The* State Transition Graph *(STG) of an ABN $B = \{f_1, \ldots, f_n\}$ over $Var_n$ is a directed graph $STG(B) = (V, E)$ where $V = \mathbb{B}^n$ (the* state space*) and $E \subseteq V \times V$ (the* transition relation*) is defined such that $(x, y) \in E$ if and only if there is a variable $v \in Var_n$ such that $y = x[v \leftarrow f_v(x)]$.*

As is customary, we write $x \to y$ when $(x, y) \in E$ and $x \to^* y$ when $(x, y) \in E^*$ (the transitive and reflexive closure of $E$). Furthermore, we can write $x \to_v y$ when the transition from $x$ to $y$ updates exactly the variable $v \in Var_n$. Note that for any $x \neq y$, either $x \not\to y$, or there is exactly one $v$ s.t. $x \to_v y$.

$STG(B)$ contains a self-loop $x \to x$ whenever $f_v(x) = x_v$. Note that some authors additionally require that the self-loops in $STG(B)$ are only present for states where no other outgoing edge is possible [2]. However, the presence of self-loops is not relevant for the problems studied in this paper, hence we use the simpler definition given above.

**Example S1.2** *The asynchronous STG of the BN from Example S1.1 is shown in Fig. S1. In state* 0000*, the BN can non-deterministically transition to states*

1000 *and* 0100*, because* $f_1(0000) = 1$ *and* $f_2(0000) = 1$*, respectively. Meanwhile,* $f_3(0000) = f_4(0000) = 0$*, hence* 0000 *also has a self-loop which is omitted in Fig. S1 by convention.*

##### S1.2.1 Restricted networks

We often need to reason about the behaviour of a network $B$ *within* a particular subspace $X \in \mathbb{B}^n_\star$. To this end, we write $B[X]$ to denote the *restriction* of $B$ to subspace $X$. Informally, $B[X]$ is a network where all fixed variables of $X$ are substituted for their corresponding constant values and removed.

This introduces the need to match variables (and states) of the "full" network $B$ to the variables of the restricted network $B[X]$ and vice versa. To simplify notation, we typically assume that it is clear from context which combination of full and restricted network is being considered. We then set $n_{[X]} = |\mathit{free}(X)|$ and define $\mathit{Var}_{n_{[X]}} = \mathit{Var}_n \setminus \mathit{fixed}(X)$ to be the set of variables that are free in $X$, i.e. the variables of $B[X]$. We can then formally define $B[X] = \{f'_{v_1}, \ldots, f'_{v_{n_{[X]}}}\}$ as a Boolean network such that $f'_v$ is constructed by applying $f_v[v' \leftarrow X_{v'}]$ for every $v' \in \mathit{fixed}(X)$ and removing (now redundant) variables from $\mathit{fixed}(X)$ ($f_v$ being the update function in the unrestricted network $B$).

For a particular state $x \in \mathcal{S}(X)$, we then use the *projection* operator $(x \downarrow X) \in \mathbb{B}^{n_{[X]}}$ to mean "state from $\mathbb{B}^{n_{[X]}}$ corresponding to $x \in \mathcal{S}(X)$ after removing values of variables fixed by $X$". Symmetrically, we define the *extension* operator $(x \uparrow X) \in \mathcal{S}(X)$ to be the full "unrestricted" state obtained by extending $x \in \mathbb{B}^{n_{[X]}}$ with all fixed values of $X$. We can define analogous projection and extension operators also for subspaces and sets of states (here, the projection and extension are applied element-wise).

Given a network $B$, subspace $X$ and the restricted network $B[X]$, we can observe the following basic property:

**Corollary S1.1** *The sub-graph of* $STG(B)$ *induced by states* $\mathcal{S}(X)$ *is isomorphic to the graph* $STG(B[X])$*.*

**Example S1.3** *Consider the four-variable network from Example S1.1 (also recall that the STG of this network is shown in Figure S1), and let us set* $X = 10{\star}{\star}$*. A restricted network* $B[X]$ *retains the last two variables, since the first two variables are fixed by* $X$*. We refer to these as* $v'_3$ *and* $v'_4$*. As per the discussion above, we get that* $f_{v'_3}(x) = 1 \vee 0 = 1$ *and* $f_{v'_4}(x) = 1 \veebar x_4 = \neg x_4$*. Clearly,* $STG(B[X])$ *generated by* $f_{v'_3}$ *and* $f_{v'_4}$ *has the same transitions as the states* $\{1000, 1001, 1010, 1011\}$ *shown in Figure S1. In particular, note that* $f_{v_4}(x_1) = f_{v'_4}(x_1 \downarrow X)$ *for all* $x_1 \in \mathcal{S}(X)$ *and* $f_{v'_4}(x_2) = f_{v_4}(x_2 \uparrow X)$ *for all* $x_2 \in \mathbb{B}^2$ *(analogously for* $v_3$ *and* $v'_3$*).*

##### S1.2.2 Network influence graph

Even though each $f_v$ can in theory depend on all network variables, this is rarely true in practice. The variable $u \in \mathit{Var}_n$ is *essential* in $f_v$ when there is a combination of inputs where the output of $f_v$ depends on the value of $u$:

$$\exists x \in \mathbb{B}^n.\ f_v(x[u \leftarrow 0]) \neq f_v(x[u \leftarrow 1])$$

The set $in(f_v) \subseteq Var_n$ denotes the set of variables essential in $f_v$. Variable $v$ is called a *network output* if it is not essential in any update function. Similarly, $v \in Var_n$ is called a *network input* (or *source*) when $f_v(x) \equiv x_v$ for all $x \in \mathbb{B}^n$. We say that $v \in Var_n$ is a *constant* when $f_v(x) \equiv true$ or $f_v(x) \equiv false$ for all $x \in \mathbb{B}^n$.

In a biological context, influences among nodes in a BN are typically also *locally monotonic*, meaning that an increase of an input value can only increase (positive monotonicity; *activation*) or decrease (negative monotonicity; *inhibition*) the function output:

$$\forall x \in \mathbb{B}^n.\ f_v(x[u \leftarrow 0]) \leq f_v(x[u \leftarrow 1]) \quad (u \text{ is a positive input of } f_v)$$
$$\forall x \in \mathbb{B}^n.\ f_v(x[u \leftarrow 0]) \geq f_v(x[u \leftarrow 1]) \quad (u \text{ is a negative input of } f_v)$$

We define $in_{\oplus}(f) \subseteq in(f)$ and $in_{\ominus}(f) \subseteq in(f)$ to be the subsets of positive and negative essential function inputs, respectively. Note that $in_{\oplus}(f)$ and $in_{\ominus}(f)$ are always disjoint. A function $f$ is locally monotonic when $in(f) = in_{\oplus}(f) \cup in_{\ominus}(f)$. Furthermore, a network is locally monotonic if all its functions are locally monotonic. Equivalently, a function is locally monotonic if and only if it can be written in disjunctive normal form where positive and negative function inputs only appear as positive and negative literals, respectively [3].

**Example S1.4** *Consider the BN $B$ from Example S1.1 (rules of this BN are $f_1(x) = \neg x_2$; $f_2(x) = \neg x_1$; $f_3(x) = x_1 \vee x_2$; $f_4(x) = x_1 \veebar x_4$). The regulatory functions $f_1$, $f_2$, and $f_3$ are locally monotonic, but $f_4$ is not. In the state* 0000*, increasing $x_4$ changes the output of $f_4$ from* 0 *to* 1 *(i.e., $x_4$ acts like a positive input) whereas in the state* 1000 *increasing $x_4$ changes the output of $f_4$ from* 1 *to* 0 *(i.e., $x_4$ acts like a negative input). Thus, $x_4$ is not an element of $in_{\oplus}(f_4)$ or $in_{\ominus}(f_4)$. A similar example can be constructed to show that $x_1$ also has this property. In contrast, $f_1$, $f_2$, and $f_3$ are locally monotonic (specifically, $in_{\ominus}(f_1) = \{x_2\}$, $in_{\ominus}(f_2) = \{x_1\}$, and $in_{\oplus}(f_3) = \{x_1, x_2\}$).*

To study the essentiality and monotonicity of a Boolean network $B$, we then consider its (signed) influence graph $IG(B)$:

**Definition S1.4** *The* influence graph *$IG(B)$ of a BN $B = \{f_1, \ldots, f_n\}$ is an edge-labelled directed graph $IG(B) = (V, E)$ with edge labeling function $\sigma : E \to \{-1, 0, 1\}$ where $V = Var_n$ is the set of Boolean variables, $E = \{(u, v) \subseteq V \times V \mid u \in in(f_v)\}$ contains an edge from $u$ to $v$ if and only if $u$ is essential in $f_v$, and the sign of each edge, given by $\sigma$, is defined via*

$$\sigma((u,v)) = \begin{cases} +1 & \text{if } u \in in_{\oplus}(f_v) \text{ (positive sign)} \\ -1 & \text{if } u \in in_{\ominus}(f_v) \text{ (negative sign)} \\ \pm 0 & \text{otherwise (ambiguous sign)} \end{cases}$$

*for $(u, v) \in E$.*

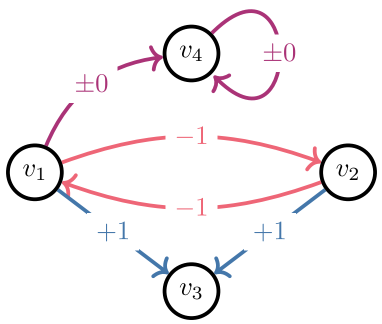


Figure S2: Influence graph of the BN from Example S1.1 (rules $f_1(x) = \neg x_2$; $f_2(x) = \neg x_1$; $f_3(x) = x_1 \vee x_2$; $f_4(x) = x_1 \veebar x_4$). Signs $+1$ (blue), $-1$ (red), and $\pm 0$ (purple) denote the positive, negative, and non-monotonic interactions, respectively.

The sign of a cycle in $IG(B)$ is given by the product of its edge signs, meaning each cycle is either positive, negative, or ambiguous. Recall that a set of nodes $F$ that intersects every cycle of $IG(B)$ is called a *Feedback Vertex Set* (FVS) of $IG(B)$. A *positive* (resp. *negative*) FVS (also denoted PFVS and NFVS) of $IG(B)$ is a set of nodes that intersects every cycle of positive or ambiguous (resp. negative or ambiguous) sign in $IG(B)$. Trivially, an FVS is also a positive FVS and a negative FVS.

A single $IG(B)$ can have many feedback vertex sets. In particular, $F = Var_n$ is always *trivially* an FVS. As such, we are typically interested in the *minimum* FVS (although such FVS is also not necessarily unique). However, finding a minimum FVS (both general and positive or negative) is known to be NP-complete [4, 5]. As such, we will often content ourselves with a sufficiently small but non-minimum FVS approximation instead.

Finally, note that the edges of an influence graph of a restricted network $B[X]$ are a subset of the edges of $IG(B)$. However, the restriction can change ambiguous signs to positive or negative, or it can eliminate edges even between variables from $free(X)$ (i.e. $IG(B[X])$ is not necessarily the full sub-graph of $IG(B)$ induced by nodes $free(X)$). As such, a minimum FVS of $IG(B)$ is also an FVS of $IG(B[X])$, but a smaller FVS can be often obtained through direct analysis of $IG(B[X])$.

**Example S1.5** *The influence graph $IG(B)$ of the network $B$ defined in Example S1.1 is shown in Fig. S2 (rules $f_1(x) = \neg x_2$; $f_2(x) = \neg x_1$; $f_3(x) = x_1 \vee x_2$; $f_4(x) = x_1 \veebar x_4$). The graph has no purely negative cycles, one purely positive cycle ($v_1 \to v_2 \to v_1$), and one ambiguous sign cycle (the self loop on $v_4$). There are two minimum feedback vertex sets among the subsets of $Var_4$, specifically, $\{v_1, v_4\}$ and $\{v_2, v_4\}$. These are also minimum positive feedback vertex sets because the self-loop on $v_4$ must be treated as both a positive and a* *negative feedback loop. The minimum NFVS is $\{v_4\}$; since the $v_1 \leftrightarrow v_2$ feedback loop has a positive sign, it need not be intersected by the NFVS.*

#### S1.3 Long-term behaviour of Boolean networks

The most important biological question concerning Boolean networks is: *What happens to the evolving state of the BN in the long-term?* In this section, we introduce mechanisms through which this question is studied.

##### S1.3.1 Boolean space percolation

Given a subspace of network states $X \in \mathbb{B}^n_\star$, the single-step percolation operator $\mathcal{P}$ acting on $X$ produces a subspace $\mathcal{P}(X)$ with fixed variables given by those of $X$ together with the free variables of $X$ whose update functions are invariant on $X$. That is,

$$(\mathcal{P}(X))_i = \begin{cases} X_i & \text{if } X_i \neq \star \\ b & \text{if } X_i = \star \text{ and } (f_i(x) = b \text{ for all } x \in \mathcal{S}(X)) \\ \star & \text{otherwise} \end{cases} \tag{4}$$

The percolation operator $\mathcal{P}^\infty$ is obtained by repeated application of the single-step percolation operator $\mathcal{P}$ until fixed-point. Note that because a Boolean network has only finitely many variables, $\mathcal{P}^\infty$ always converges and there is some (finite) integer $k \leq n$ for which $\mathcal{P}^k(X) = \mathcal{P}^{k+1}(X) = \mathcal{P}^\infty(X)$.

**Example S1.6** *Consider the BN from Example S1.1 (rules $f_1(x) = \neg x_2$; $f_2(x) = \neg x_1$; $f_3(x) = x_1 \vee x_2$; $f_4(x) = x_1 \veebar x_4$) and the Boolean subspace $\star 1 \star\star$. For this space, we have $\mathcal{P}(\star 1 \star\star) = \star 1 1 \star$, because for every $x \in \star 1 \star\star$, we have $f_3(x) = x_1 \vee true = true$. Similarly, $\mathcal{P}(\star 11 \star) = 011\star$, because for every $x \in \star 11 \star$, we have $f_1(x) = \neg true = false$. The space $011\star$ cannot be further percolated because the only remaining free variable, $x_4$, has an update function that simplifies to $f_4(x) = x_4$ for $x \in 011\star$. This is neither a tautology nor a contradiction. Thus, $\mathcal{P}^\infty(\star 1 \star\star) = \mathcal{P}^2(\star 1 \star\star) = 011\star$ in this BN.*

##### S1.3.2 Trap sets, trap spaces and attractors

Next, consider an arbitrary set of states $T \subseteq \mathbb{B}^n$ of BN $B$. Such $T$ is a *trap set* if and only if $T$ cannot be escaped within $STG(B)$. That is, for every $x \in T$ we have that $x \to y$ implies $y \in T$. Such a set can be also called *forward closed*, as it is closed under the transition relation of $STG(B)$.

Furthermore, the *inclusion minimal* trap sets of $STG(B)$ are called *attractors* and we write $\mathcal{A}(B)$ to denote the set of all attractors of BN $B$. Typically, we distinguish between *fixed-point* (a single state), *cyclic* (a single cycle), and *complex* (arbitrary set) attractors. Attractors are crucial when studying the long-term behaviour of Boolean networks, as any *fair* run within $STG(B)$ is eventually trapped in an attractor. Alternatively, attractors can be also defined as the bottom (terminal) strongly connected components of $STG(B)$. However, note that the exact attractors depend on the choice of the update scheme. Here, $\mathcal{A}(B)$ denotes the attractors in the *asynchronous* update, but we could similarly obtain a different set of attractors under the synchronous (or other) update.

Another special class of trap sets is called *trap spaces*. A trap space is a trap set $T$, where $T$ is a Boolean subspace (i.e. $T$ corresponds to one of the members of $\mathbb{B}^n_\star$). We write $\mathcal{T}_{all}(B)$ to denote the set of all trap spaces of a BN $B$. Note that a Boolean network has the same trap spaces regardless of the network's update scheme [6]. Similar to attractors, we can define the *minimal trap spaces* as the inclusion-minimal Boolean spaces that are also trap sets. $\mathcal{T}_{min}(B)$ denotes the set of such trap spaces for a BN $B$.

However, as opposed to an attractor, a minimal trap space is not necessarily a strongly connected component of $STG(B)$. Nevertheless, every minimal trap space contains *at least* one attractor, regardless of the update scheme (this follows from the definition of an attractor as an inclusion minimal trap set). Hence minimal trap spaces are often used to approximate asynchronous attractors [7, 8]. However, not *every* attractor is necessarily located in a minimal trap space. As such, finding efficient methods to *a priori* assess the quality of this approximation remains an open problem. In our text, we write $\mathcal{T}(A)$ to denote the smallest trap space in which an attractor $A \in \mathcal{A}(B)$ resides. As per previous discussion, often we have $\mathcal{T}(A) \in \mathcal{T}_{min}(B)$, but this is not a rule.

A special case where trap spaces and attractors fully overlap are the so-called fixed-points, or single state attractors. Every fixed-point is guaranteed to be both an attractor and a minimal trap space. The set of fixed-points of a network $B$ is denoted $\mathcal{T}_{fix}(B)$.

Symmetrically to the minimal trap space, we can define a *maximal* trap space $T$ within an arbitrary subspace $X \in \mathbb{B}^n_\star$. We say that $T$ is a maximal trap space within $X$ if it is an inclusion-maximal proper subspace of $X$ (i.e. $T \subset X$) that is also a trap space. We then write $\mathcal{T}_{max}(X, B) \subseteq \mathbb{B}^n_\star$ to denote the set of all maximal trap spaces of $B$ within $X$. Note that when $X = \star^n$, $\mathcal{T}_{max}(X, B)$ is the set of all globally maximal trap spaces of $B$ as defined in [6]. We denote this $\mathcal{T}_{max}(B) = \mathcal{T}_{max}(\star^n, B)$.

Finally, we briefly discuss the impact of network restriction on trap sets and trap spaces. Clearly, if $X \in \mathbb{B}^n_\star$ is a trap space, then all attractors of $B$ that are present in $\mathcal{S}(X)$ are also attractors of $B[X]$ (after projection), and $B[X]$ has no other attractors. A slightly more general conclusion can be also inferred for trap spaces: Any $Y$ that is a subspace of $X$ is a trap space of $B$ if and only if its projection is a trap space of $B[X]$.

**Example S1.7** *Let us again consider the Boolean network $B$ from Example S1.1 and recall that Fig. S1 depicts the asynchronous $STG(B)$. The network has two single state attractors,* 0110 *(red) and* 0111 *(green). Furthermore, there is a single cyclic attractor* $\{1010, 1011\}$ *(blue). Coincidentally, these also correspond to the minimal trap spaces of the whole network:* 0110*,* 0111*, and* $101\star$*.*

*There are six additional non-minimal trap spaces:*

$$\{\ 01\star0, 01\star1, 011\star, 01\star\star, 10\star\star, \star\star\star\star\ \}.$$

*However, all of these except for $\star\star\star\star$ and $011\star$ percolate to a smaller trap space. Subspace $01\star0$ percolates to $0110$, $01\star1$ percolates to $0111$, $01\star\star$ percolates to $011\star$, and $10\star\star$ percolates to $101\star$. Consequently, the globally maximal trap spaces (i.e. maximal within $\star\star\star\star$) are $\{01\star\star, 10\star\star\}$.*

*We see that the network is first presented with a non-deterministic choice of committing to either $01\star\star$ or $10\star\star$. In the first case, another non-deterministic choice leads to one of the single state attractors. In the second case, the network converges to the cyclic attractor.*

##### S1.3.3 Succession diagrams

Example S1.7 hints at a possible hierarchy of trap spaces based on inclusion. We formalise this hierarchy using the notion of a *succession diagram*, which was first introduced in [7].

Originally, succession diagrams use the concept of *stable motifs* to derive the hierarchy of trap spaces [7, 9]. Here, each stable motif represents a possible decision leading to a successively more restricted nested trap spaces. However, it has been shown that in this application, the stable motifs are in-fact equivalent to maximal trap spaces [9]. As such, we present a more succinct definition of the succession diagram in terms of maximal trap spaces and percolation that is based on the definition given in [9] but adapted to our notation[1]:

**Definition S1.5** *The* succession diagram $SD(B) = (r, V, E)$ *of a Boolean network $B$ is a rooted directed acyclic graph such that:*

- *The vertices of this graph are the percolated trap spaces of $B$:*

$$V = \{\ T \in \mathbb{B}^n_\star \mid \exists T' \in \mathcal{T}_{all}(B).\ T = \mathcal{P}^\infty(T')\ \} \tag{5}$$

- *The root is the percolation of the universal trap space, i.e., $r = \mathcal{P}^\infty(\star^n)$.*
- *An edge between trap spaces $T$ to $T'$ (i.e., $(T, T') \in E$) exists if and only if $T' = \mathcal{P}^\infty(U)$ for some $U \in \mathcal{T}_{max}(T, B)$.*

Intuitively, the succession diagram follows the inclusion hierarchy of trap spaces, but "skips ahead" through any trivial relationships which can be easily deduced through percolation. Note that the leaf (i.e., terminal) nodes of $SD(B)$ are exactly the minimal trap spaces $\mathcal{T}_{min}(B)$ [9].

[1] In [7] and [9], the succession diagram is defined in terms of the prime implicant structure of the update functions using an *expanded network* or *prime implicant hypergraph* rather than directly from the trap spaces. The succession diagram defined in [9] is essentially the line graph of the succession diagram defined in [7]. Our definition more closely follows [9]. In contrast to the definition given in [9], we describe the nodes of the succession diagram using the percolated trap spaces, whereas [9] uses the unpercolated trap spaces, treating the percolation as implicit. When multiple maximal trap spaces give rise to the same trap space under percolation, our definition merges these maximal trap spaces into a single node, whereas [9] treats them as separate nodes.

The succession diagram is useful for studying the control and decision making in the associated Boolean network [10, 11]. Of particular note is that the resulting control interventions are guaranteed to work in any update scheme. Note that, in [9], the succession diagram is also used to identify all attractors of a Boolean network under the asynchronous update scheme.

As argued in [9] and depicted in Figure S3, nodes of the succession diagram may be thought of as decision points in the dynamics of the network, analogous to peaks in the epigenetic landscape of Waddington [12].

**Example S1.8** *Recall the Boolean network $B$ from Example S1.1 and the discussion about its trap spaces in Example S1.7. Clearly, the succession diagram $SD(B) = (r, V, E)$ of this network consists of five trap spaces (with $r = \star\star\star\star$):*

$$V = \{\ \star\star\star\star, 101\star, 011\star, 0111, 0110\ \}$$

*There are four edges: First, from* $\star\star\star\star$ *to* $101\star$ *and* $011\star$*, because* $\mathcal{P}^\infty(10\star\star) = 101\star$ *and* $\mathcal{P}^\infty(01\star\star) = 011\star$*. Second, from* $011\star$ *to* $0110$ *and* $0111$*. Trap spaces* $01\star0$ *and* $01\star1$ *do not directly appear in the succession diagram because they are among the maximal subspaces of* $01\star\star$*, which percolates to* $011\star$*. Consequently, these subspaces also percolate to* $0110$ *and* $0111$*, respectively.*

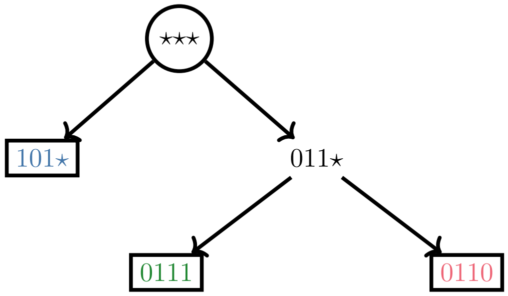


Figure S3: Succession diagram of the Boolean network from Example S1.1 (rules $f_1(x) = \neg x_2$; $f_2(x) = \neg x_1$; $f_3(x) = x_1 \vee x_2$; $f_4(x) = x_1 \veebar x_4$). Leaf node labels are colored to coincide with the state color of attractors in the STG of Fig. S1.

#### S1.4 Petri nets

Finally, we recall the notion of *Petri nets*. Petri nets are widely used in biological modelling [13, 14, 15, 16, 17, 18] and they have also proven useful in the characterisation of long-term behaviour of Boolean networks [19, 20, 21, 22], including for the identification of trap spaces and attractors.

**Definition S1.6** *A* Petri Net *(PN) is a weighted directed bipartite graph $PN = (P, T, E, W)$, where $P$ and $T$ are non-empty, finite and disjoint sets of vertices called* places *and* transitions*, respectively, and where $W : E \to \mathbb{N}$ is the weight function that assigns a positive integer weight to every arc leading from a place to a transition (or vice versa).*

Additionally, a *marking* of a Petri net is a mapping $M \colon P \to \mathbb{N}$ that assigns a number of *tokens* to each place. A place $p \in P$ is *marked* by $M$ if and only if $M(p) > 0$. We write $\mathit{pred}(x)$ (resp. $\mathit{succ}(x)$) to represent the set of PN vertices that have a non-zero weighted arc leading to (resp. coming from) vertex $x$.

In this work, we are only interested in a specific class of Petri nets called *one-safe* (or 1-safe) PNs. A Petri net is considered one-safe if every admissible marking $M$ only marks each place with up to one token, that is $M(p) \leq 1$ for all $p \in P$.

Every such Petri net can be described using only arcs with weights 0 and 1. However, this simple requirement is not always sufficient: There are Petri nets with only 0/1 arcs that are not one-safe. Nevertheless, in our case it will be easy to show that the considered Petri nets are always one-safe by construction. Finally, in a one-safe Petri net, we do not have to explicitly write arc weights, since each arc is either shown (weight 1) or not (weight 0). Similarly, each mapping $M$ can be simplified to a subset of marked places (i.e., $M \subseteq P$).

Now, we can formalise the dynamics of such one-safe Petri net. First, a transition $t \in T$ is considered *enabled* in marking $M$ iff $\mathit{pred}(t) \subseteq M$. The *firing* of $t$ then leads to a new marking $M'$ such that $M' = (M \setminus \mathit{pred}(t)) \cup \mathit{succ}(t)$. In other words, the firing of the transition $t$ removes a token from every $p \in \mathit{pred}(t)$ and places a token into all $p' \in \mathit{succ}(t)$. When multiple transitions are enabled, we typically consider that each can be fired non-deterministically [23], similar to the asynchronous update described for Boolean networks.

##### S1.4.1 Encoding Boolean dynamics

The link between Boolean networks and one-safe Petri nets was originally established in [15] to enable analysis of BNs through formal methods like model-checking. The basic Boolean encoding was later extended to multi-valued logical models in [14, 21].

Since the Boolean encoding is crucial to this paper, we briefly recall it here. Consider a BN $B = \{f_1, \ldots, f_n\}$. We construct a corresponding $PN(B)$ such that for each variable of $B$, we create two corresponding places $P = \bigcup_{v \in \mathit{Var}_n} \{n_v, p_v\}$. Intuitively, when marked, $p_v$ represents that variable $v$ is active (i.e. 1), while marked $n_v$ represents inactivity (i.e. 0).

Now, assume that we have a disjunctive normal form (DNF) representation of $(f_v(x) \wedge \neg x_v)$ for each $v \in \mathit{Var}_n$. Each clause $c$ of the $(f_v(x) \wedge \neg x_v)$ DNF formula then generates one $PN(B)$ transition $t_c \in T$ which moves a token from place $n_v$ to the place $p_v$ when the clause $c$ is satisfiable by the current PN marking. Specifically, we always have $n_v \in \mathit{pred}(t_c)$ and $p_v \in \mathit{succ}(t_c)$ for the network variable $v$. For any remaining variable $u \neq v$ that appears in the clause $c$ as a *positive* literal, we have that $p_u \in \mathit{pred}(t_c)$ and $p_u \in \mathit{succ}(t_c)$. Analogously, for variables that appear as negative literals, we use the place $n_u$ instead of $p_u$. Note that no variable can appear as both positive and negative, as such clause is clearly unsatisfiable. Finally, a symmetrical process generates the transitions from $p_v$ to $n_v$ using the DNF of $(\neg f_v(x) \wedge x_v)$ instead of $(f_v(x) \wedge \neg x_v)$.

Intuitively, the role of each transition is to check whether the associated

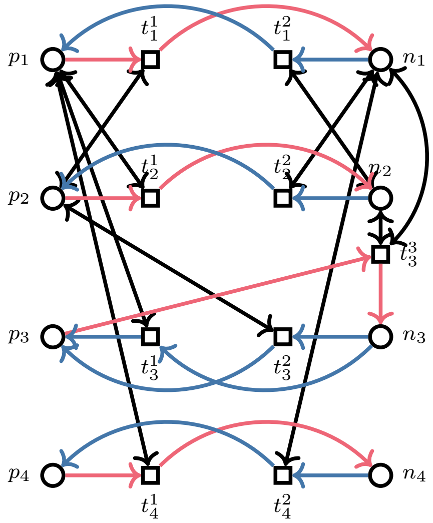


Figure S4: Petri net encoding of the Boolean network from Example S1.1 (rules $f_1(x) = \neg x_2$; $f_2(x) = \neg x_1$; $f_3(x) = x_1 \vee x_2$; $f_4(x) = x_1 \veebar x_4$). Circles and squares denote places and transitions, respectively. For each transition, the (directional) arcs representing the variable update are shown as either blue (increase) or red (decrease). The remaining bi-directional arcs ensuring the satisfiability of the DNF clause are shown as black.

clause is satisfied by taking a token from $p_u$ or $n_u$ for all relevant $u \neq v$ and then returning it back to its original place. If this condition succeeds and the transition is fire-able, we can safely move the token from $n_v$ to $p_v$ for the variable that is actually being updated.

**Example S1.9** *Let us consider the BN $B$ from Example S1.1. By using the process described above, we get the following DNFs:*

$$
\begin{aligned}
&f_1(x) \wedge \neg x_1 \equiv \neg x_2 \wedge \neg x_1 && \neg f_1(x) \wedge x_1 \equiv x_2 \wedge x_1 \\
&f_2(x) \wedge \neg x_2 \equiv \neg x_1 \wedge \neg x_2 && \neg f_2(x) \wedge x_2 \equiv x_1 \wedge x_2 \\
&f_3(x) \wedge \neg x_3 \equiv (x_1 \wedge \neg x_3) \vee (x_2 \wedge \neg x_3) && \neg f_3(x) \wedge x_3 \equiv \neg x_1 \wedge \neg x_2 \wedge x_3 \\
&f_4(x) \wedge \neg x_4 \equiv x_1 && \neg f_4(x) \wedge x_4 \equiv \neg x_1
\end{aligned}
$$

*The Petri net constructed based on these DNF representations is shown in Figure S4. Note that in this simple example, every combination except for $(f_3(x) \wedge \neg x_3)$ generates a DNF with a single clause, and consequently a single transition.*

Formally, we can establish a relationship between Boolean network $B$ and its Petri net encoding $PN(B)$ as follows: First, we observe that for every state $x \in \mathbb{B}^n$, there is a marking $M_x$ which encodes $x$ in terms of the $PN(B)$ places.

Specifically, for every $v \in Var_n$, we have $(p_v \in M_x) \Leftrightarrow x_v = 1$ and $(n_v \in M_x) \Leftrightarrow x_v = 0$. For example, a state 101 is encoded as a marking $\{p_1, n_2, p_3\}$. As a result, we always have $M_x(p_v) + M_x(n_v) = 1$.

Subsequently, we have that the Petri net $PN(B)$ can transition from marking $M_x$ to marking $M_y$ if and only if the corresponding $STG(B)$ has a transition $x \to y$. That is, the encoding is completely faithful with respect to the state-transition graph of $B$ and thus preserves the dynamics of the original BN [24]. Finally, assuming $M(p_v)+M(n_v) = 1$, such Petri net is clearly one-safe, as each transition moves a single token between $p_v$ and $n_v$.

**Encoding construction** Note that transforming a Boolean network into its Petri net encoding relies on obtaining conditions for the activation and deactivation of the individual variables, here represented by the DNF of $(f_v(x) \wedge \neg x_v)$, resp. $(\neg f_v(x) \wedge x_v)$. In [15], this is based on the complete truth table of each update function. But as shown in Appendix 1 of [21], any DNF is in fact sufficient. In particular, there is no requirement for this DNF to be based on prime implicants, nor is it required that all implicants are covered (as is the case for the Blake canonical form used in [25, 11]). As such, `biobalm` implements the DNF transformation by encoding the function into a BDD and enumerating the satisfiable clauses of this BDD, as proposed in [19].

##### S1.4.2 Siphons and traps

Siphons or traps are static and classical properties of Petri nets [26]. Note however that the use of siphons or traps for the analysis of biological models, though it is not new, has been mostly relevant to the ODE-based continuous semantics of chemical reaction networks [16, 27, 28]. We recall here the basic definitions establishing that to produce (resp. consume) something in a siphon (resp. trap) you must consume (resp. produce) something from the siphon (resp. trap). This corresponds to the idea that a siphon (resp. trap) is a set of places that once unmarked (resp. marked) remains unmarked (resp. marked).

**Definition S1.7** *A* siphon *of a Petri net* $(P, T, W)$ *is a set of places* $S$ *such that:*

$$\forall t \in T, S \cap succ(t) \neq \emptyset \Rightarrow S \cap pred(t) \neq \emptyset.$$

*Note that* $\emptyset$ *is trivially a siphon.*

**Definition S1.8** *A* trap *of a Petri net* $(P, T, W)$ *is a set of places* $S$ *such that:*

$$\forall t \in T, S \cap pred(t) \neq \emptyset \Rightarrow S \cap succ(t) \neq \emptyset.$$

*Note that* $\emptyset$ *is trivially a trap.*

Let $pred(S) := \bigcup_{s \in S} pred(s)$ and $succ(S) := \bigcup_{s \in S} succ(s)$. If $S = \emptyset$, then conventionally $pred(S) = succ(S) = \emptyset$. We have two important properties on siphons and traps [29] as follows.

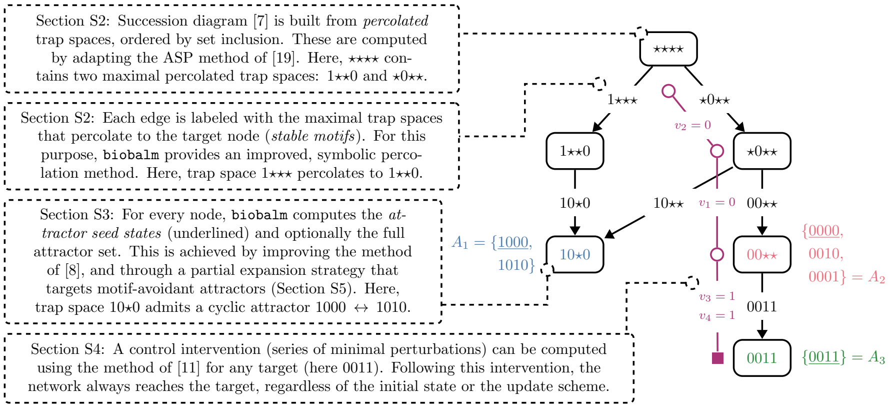


Figure S5: Summary of the `biobalm` features introduced in the main paper. The diagram is annotated with the network attractor sets and a chosen control intervention. The update functions and the STG of the network represented in this diagram are available in Section S8.1.

**Proposition S1.1** *A set $S$ of places is a siphon of a Petri net $(P, T, W)$ if and only if $pred(S) \subseteq succ(S)$.*

**Proposition S1.2** *A set $S$ of places is a trap of a Petri net $(P, T, W)$ if and only if $succ(S) \subseteq pred(S)$.*

Let $PN$ be a one-safe Petri net. Its *complementary* Petri net (say $PN'$) is the Petri net that shares the same sets of places and transitions with $PN$, but all the arcs are reversed. It is known that a siphon (resp. trap) of $PN$ is a trap (resp. siphon) of $PN'$ [29].

**Example S1.10** *Let us consider the Petri net shown in Figure S4. Set $S_1 = \{p_1, p_2\}$ is not a siphon but a trap because $pred(S_1) = \{t_1^1, t_1^2, t_2^1, t_2^2, t_3^1, t_3^2, t_4^1\}$ and $succ(S_1) = \{t_1^1, t_2^1, t_3^1, t_3^2, t_4^1\}$ (i.e., $succ(S_1) \subseteq pred(S_1)$). $S_2 = \{n_1, p_2, n_3\}$ is a siphon but not a trap because $pred(S_2) = \{t_1^1, t_2^2, t_3^2, t_3^3, t_4^2\}$ and $succ(S_2) = \{t_1^1, t_1^2, t_2^1, t_2^2, t_3^1, t_3^2, t_3^3, t_4^2\}$ (i.e., $pred(S_2) \subseteq succ(S_2)$).*

### S2 Succession diagram construction

This section outlines in detail the individual methods used for the construction of succession diagrams in the tool `biobalm`. In particular, this covers trap space computation, percolation, and the lazy expansion of SD nodes. Visually, the features of the succession diagram are summarized in Figure S5.

#### S2.1 Trap space computation

The employed trap space detection method is based on the one presented in [19]. Specifically, it relies on the Petri net encoding of Boolean dynamics as presented in Section S1.4.1 and Theorems 1 and 2 of [19], which link the maximal (conflict-free) siphons of the Petri net encoding $PN$ with the minimal trap spaces of a Boolean network $B$. However, we extend the method in several non-trivial aspects to make it viable for the construction of succession diagrams.

##### S2.1.1 Optimizing the Petri net encoding

The original package `trappist` from [19] uses a binary decision diagram data structure from the package `PyEDA`[2] to generate the disjunctive normal form necessary to create the Petri net encoding $PN(B)$. The size of this DNF (i.e. the number of clauses) is crucial to the performance of the method. BDDs can be used to compute the minimal DNF (the set of essential prime implicants) [30], but it is still non-trivial to obtain and depends on the BDD variable ordering.

Instead, we propose a simple heuristic method to optimize the number of DNF clauses in reasonably small functions, described in Algorithm S1. Here, $\textsc{vars}(f)$ denotes the subset of $Var_n$ that is essential in $f$ and $|f|$ is the size of the function in terms of BDD node count. Recall that $f[x \leftarrow b]$ denotes the substitution of variable $x$ for constant $b$ in the function $f$. The method greedily selects a variable that decomposes the BDD into smallest sub-products and then uses this variable to recursively compute the DNF. Note that compared to naive DNF construction, this method is less sensitive to variable ordering, since each recursive branch can greedily select different variables. However, it also involves greater number of BDD operations, as the function is recursively subdivided into smaller BDDs.

**def** $\textsc{dnf}(f : \texttt{BDD})$**:**
 **if** $f$ ***is*** *false* **then return** $\{\}$;
 **if** $f$ ***is*** *true* **then return** $\{\{\}\}$;
 $v \leftarrow v \in \textsc{vars}(f)$ with minimal $|f[v \leftarrow 0]| + |f[v \leftarrow 1]|$;
 $d_0, d_1 \leftarrow \textsc{dnf}(f[v \leftarrow 0]), \textsc{dnf}(f[v \leftarrow 1])$;
 **return** $\{\ c \cup \{v\} \mid c \in d_1\ \} \cup \{\ c \cup \{\neg v\} \mid c \in d_0\ \}$;

**Algorithm S1:** Heuristic algorithm for reducing DNF size.

##### S2.1.2 Maximal trap spaces with PN encoding restriction

Originally, [19] only describes a method for computing minimal trap spaces. Later, [31] demonstrates that the globally maximal case is symmetric (i.e., minimal siphons are used instead of maximal ones). As such, detection of globally maximal trap spaces (i.e. $\mathcal{T}_{max}(B)$) can be implemented using the subset-minimal solutions feature in the `clingo` solver (instead of subset-maximal solutions that are used to compute maximal siphons, i.e. the minimal trap spaces).

Furthermore, to generate a succession diagram, we need to repeatedly compute the set of maximal trap spaces $\mathcal{T}_{max}(X, B)$ for various $X \in \mathcal{T}_{all}(B)$. However, we only discussed the computation of $\mathcal{T}_{max}(B)$ so far.

[2] `https://pyeda.readthedocs.io/`

To compute $\mathcal{T}_{max}(X, B)$, we rely on the Petri net encoding of the restricted network $B[X]$, denoted $PN(B[X])$. In $PN(B[X])$ (resp. $B[X]$), all $fixed(X)$ variables are removed, leaving only the relevant $free(X)$ variables. Subsequently, once we obtain the globally maximal trap spaces of $B[X]$ through such $PN(B[X])$ (using `clingo` as discussed above), we can extend them to the original network $B$ by adding the missing fixed values (recall the $\uparrow$ operator), resulting in the set $\mathcal{T}_{max}(X, B)$.

Finally, note that a different $PN(B[X])$ is required for each succession diagram node (i.e. a percolated trap space $X$). To avoid repeating the relatively costly PN encoding step for each $B[X]$ (due to the DNF optimization), we propose a simple process for obtaining $PN(B[X])$ directly from the "global" $PN(B)$ encoding.

To construct $PN(B[X])$, we consider every variable $v$ that is fixed within $X$, and apply the following (w.l.o.g. assume $X_v = 1$, the other case is symmetric):

- We remove any PN transition that takes a token from $p_v$ and places it in $n_v$, or vice versa. These transitions never trigger within $B[X]$, since we know $v$ is set to a constant and removed in $B[X]$.
- At this point, all transitions that take a token from either $p_v$ or $n_v$ also place a token in the same place. I.e., the "value" of $p_v$ and $n_v$ is constant.
- We disconnect the the place $p_v$ from all the remaining transitions and remove it from the Petri net. Since $X_v = 1$, we assume this place is always marked and can be thus safely ignored.
- We remove the place $n_v$ *together* with any transitions connected to it. Since $X_v = 1$, we assume this place is never marked and thus none of the connected transitions can fire.

We denote this operation as $PN(B[X]) \leftarrow \textsc{Restrict}(PN(B), X)$ in pseudocode. Subsequently, we propose the following corollary:

**Corollary S2.1** *Let $X$ be a trap space of $B$. The globally maximal trap spaces $\mathcal{T}_{max}(B[X])$ of $B[X]$ (here computed through $PN(B[X])$ using the PN siphons method) extended with fixed variables from $X$ are equivalent to the locally maximal trap spaces $\mathcal{T}_{max}(X, B)$:*

$$\mathcal{T}_{max}(X, B) = \{T \uparrow X \mid T \in \mathcal{T}_{max}(B[X])\} \tag{6}$$

This claim is easily explained by Corollary S1.1 and the fact that $X$ is a trap space. Since $X$ is a trap space, there are no outgoing transitions from $\mathcal{S}(X)$ within $STG(B)$. Therefore, trap spaces present in $B$ within $X$ are exactly the same as trap spaces of $B[X]$ since the two graphs are isomorphic, only the variables that are constant in $X$ are removed.

Finally, let us reiterate that with this method, it is sufficient to only create the full Petri net encoding $PN(B)$ of $B$ once, and then restrict the Petri net graph directly for every computation of $\mathcal{T}_{max}(X, B)$. This means that we do not need to repeat the relatively expensive BDD-based translation from $B[X]$ into $PN(B[X])$ repeatedly.

#### S2.2 Percolation

The percolation operator $\mathcal{P}^\infty$ is an integral part of our implementation, and especially in larger networks can play a detrimental role in the tool's performance when generating a succession diagram. In `biobalm`, we implement $\mathcal{P}^\infty$ using the BDD representation of the update functions and the restriction operation.

```
def PERCOLATE(X ∈ 𝔹ⁿ_⋆):
    continue ← true;
    bdd_v ← BDD(f_v) for all v ∈ free(X);
    while continue do
        continue ← false;
        for v ∈ free(X) do
            for u ∈ VARS(bdd_v) s.t. u ∈ fixed(X) do
                bdd_v ← bdd_v[u ← X_u];
            if bdd_v ≡ 1 ∨ bdd_v ≡ 0 then
                continue ← true;
                X ← X[v ← bdd_v];
    return X;
```

**Algorithm S2:** Algorithm implementing the $\mathcal{P}^\infty$ operator.

The used percolation method is outlined in Algorithm S2. Here, we initially instantiate each relevant $f_v$ of $B$ as a BDD $bdd_v$, and subsequently proceed to restrict the BDD until it reduces to a constant, or a fixed-point is reached. Observe that the value of $bdd_v$ is preserved between iterations, and each restriction $bdd_v[u \leftarrow X_u]$ completely eliminates variable $u$ from $\textsc{vars}(bdd_v)$. Hence, every relevant restriction is applied exactly once.

#### S2.3 Lazy succession diagram construction

In `biobalm`, we propose a `SuccessionDiagram` data structure which maintains the succession diagram graph as described in Definition S1.5. Compared to Definition S1.5, we construct the succession diagram on-the-fly, meaning new nodes and edges are added dynamically as the graph is explored. This allows us to represent (and more importantly, compute) only a portion of the succession diagram when said portion is sufficient for the particular problem at hand. Still, in the following, we write "succession diagram" to mean the full succession diagram (in line with Definition S1.5), unless explicitly stated otherwise.

We consider a trap space $X$ (i.e., a vertex of the succession diagram $SD(B)$) to be *expanded* if and only if all its outgoing edges to the percolated maximal trap spaces of $\mathcal{T}_{max}(X, B)$ are also present in $SD(B)$. Any other node is *not expanded* and must have no outgoing edges.

Here, note that a minimal trap space $X$ has no outgoing edges. As such, it is technically expanded by definition. However, in our implementation, we must distinguish between cases where $X$ is proven to be minimal (and has no successors), and where $X$ is simply not expanded yet (and further successors could be discovered). We thus maintain an `expanded`$(X)$ flag for each node and update this flag once the node's successors are computed.

```
def EXPAND(X : vertex of SD(B)):
    if expanded(X) then return;
    PN(B[X]) ← RESTRICT(PN(B), X);
    for Y ∈ T_max(X, B) using PN(B[X]) do
        Y_p ← P^∞(Y);
        V(SD(B)) ← V(SD(B)) ∪ {Y_p};
        E(SD(B)) ← E(SD(B)) ∪ {(X, Y_p)};
        StableMotifs(X, Y_p) ← StableMotifs(X, Y_p) ∪ Y;
    expanded(X) ← true;
```

**Algorithm S3:** Algorithm for expansion of a single succession diagram node.

The process of expanding a single succession diagram node is outlined in Algorithm S3. Here, $SD(B)$ and $PN(B)$ are global variables and we write $V(SD(B))$ and $E(SD(B))$ to denote the vertices and edges of $SD(B)$, respectively. Finally, the Petri net $PN(B[X])$, the set $\mathcal{T}_{max}(X, B)$, and the percolation operator $P^\infty$ are computed according to the algorithms outlined in Section S2.1 and Section S2.2.

Note that for each edge of the succession diagram, we also maintain a set `StableMotifs`$(X, Y_p)$ of stable motifs which enable said edge (i.e., the maximal trap spaces of $X$ which percolate to $Y_p$). This set is typically a singleton (i.e., each edge corresponds to a unique stable motif). However, in general, more than one maximal trap space can percolate to the same subspace, meaning that the same edge can be "rediscovered" using more than one maximal trap space.

### S3 Attractor detection

In this text, we consider two flavours of the attractor detection problem:

**Problem S3.1 (Attractor sets)** *Given a BN $B$, the goal of the* attractor set problem *is to fully compute the sets $\mathcal{A}(B) = \{A_1, \dots, A_k\}$, where $A_i \subseteq \mathbb{B}^n$ are the network attractors.*

**Problem S3.2 (Attractor seeeds)** *Given a BN $B$, the goal of the* attractor seed problem *is to compute the states $\mathcal{A}_{seed}(B) = \{a_1, \dots, k_k\}$, such that there is exactly one $a_i \in \mathcal{A}_{seed}(B)$ for every $A_i \in \mathcal{A}(B)$ that satisfies $a_i \in A_i$.*

The rational behind Problem S3.2 is that in many instances, the actual attractor sets are too large (or complicated) to output fully. In such cases, it is typically useful to at least obtain a representing state for each attractor set, which is formalized by Problem S3.2. Give such attractor seed $a_i$, we can still attempt to compute the full attractor set $A_i$ if desired using any suitable reachability method, since $A_i$ is exactly the set of states reachable from $a_i$. As such, in `biobalm`, we primarily focus on Problem S3.2, but we incorporate a symbolic reachability method that can be used to solve Problem S3.1 when the attractor seed states are already known.

In `biobalm`, we use the structure of the succession diagram to propose a new, accelerated attractor detection method. The core of the method relies on the observation that each attractor is fully contained in one of the network's percolated trap spaces (i.e., succession diagram vertices), and that each minimal trap space must contain at least one attractor. Consequently, given a succession diagram, we process each of its vertices independently. With enough care, it is even unnecessary to fully expand the succession diagram during attractor search, as we show in the next section.

For each succession diagram node, the attractor detection is then based on [8], where the network's (negative) feedback vertex set is used to identify a set of candidate states that is guaranteed to cover every network attractor. In [8], this candidate set is then reduced through simulation, static analysis and explicit model checking until a single state remains for every network attractor.

We improve on this method in three major ways: First, we propose a new heuristic that helps to reduce the size of the initial candidate set with the help of an ASP solver. Second, we improve the performance of the simulation method, allowing it to opportunistically explore longer state sequences in networks where it is the most effective. Finally, we replace the explicit model checking step with symbolic analysis using AEON [32], which appears to be more effective for larger networks and can be used to compute the full set of attractor states, not just a single seed state (however, this step is by default optional).

#### S3.1 Attractor landscape of a succession diagram

First, we observe a key lemma which relates network attractors to succession diagram nodes and enables our attractor detection method. In the following, we consider $\textsc{MotifStates}(X)$ to be the set of network states that correspond to all the stable motifs (unpercolated maximal trap spaces) within $X$:

$$\textsc{MotifStates}(X, B) = \bigcup_{T \in \mathcal{T}_{max}(X,B)} \mathcal{S}(T) \tag{7}$$

Note that in a fully expanded succession diagram, $\textsc{MotifStates}(X, B) = \emptyset$ if and only if $X$ is a minimal trap space.

**Lemma S3.1** *Let $B$ be a BN and let $A \in \mathcal{A}(B)$ be one of its attractors. Then for every trap space $X \in \mathcal{T}_{all}(B)$, either $A \subseteq \mathcal{S}(X)$, or $A \cap \mathcal{S}(X) = \emptyset$.*

**Proof** This is a well known property that follows from the fact that every trap space $X$ is a trap set, meaning that there are no transitions within $STG(B)$ leading *outside* of $\mathcal{S}(X)$. Hence, not just every attractor, but every cycle within $STG(B)$ cannot cross the boundary of a trap set. □

**Lemma S3.2** *Let $B$ be a Boolean network and let $A \in \mathcal{A}(B)$ be one of its attractors. Then there exists* exactly *one trap space $X \in V(SD(B))$ such that $A \subseteq (\mathcal{S}(X) \setminus \textsc{MotifStates}(X, B))$, and this trap space is $\mathcal{T}(A)$.*

**Proof** With Lemma S3.1 in mind, it is easy to also show this slightly stronger property. Since $A$ cannot cross the boundary between two trap spaces, there must be a single trap space $X$, such that $A \subseteq \mathcal{S}(X)$ and $A \cap \mathcal{S}(Y) = \emptyset$ for every $Y \in \mathcal{T}_{max}(X, B)$. Such $X$ is the minimal trap space that still contains $A$, which we previously denoted $\mathcal{T}(A)$. Furthermore, any such $\mathcal{T}(A)$ must always be a fully percolated trap space (i.e., it is a succession diagram node). Assuming $\mathcal{P}^{\infty}(X) = Y$ s.t. $Y \neq X$, then every state within $\mathcal{S}(X)$ can eventually reach a state in $\mathcal{S}(Y)$ simply by updating the percolated variables (and by the nature of percolation, these transitions do not have a reverse counterpart anywhere in $X$), and thus it cannot be an attractor state. Consequently, for every attractor $A$, we get that $A \subseteq (\mathcal{S}(X) \setminus \textsc{MotifStates}(X, B))$ for $X = \mathcal{T}(X)$. $\square$

Note that in practice, for most attractors, this unique trap space is also one of the network's minimal trap spaces. If $A$ is a fixed-point attractor (that is, $A = \{x\}$ for some $x \in \mathbb{B}^n$), this is in fact guaranteed. However, the so called motif-avoidant attractors [9] that reside in non-minimal trap spaces are still possible and need to be accounted for.

From these two lemmas, it is clear that any attractor detection method that is applied only to the individual $\mathcal{S}(X) \setminus \textsc{MotifStates}(X)$ sets in isolation for every $X \in V(SD(B))$ still completely detects the set $\mathcal{A}(B)$. We can further simplify the problem by observing the following:

**Lemma S3.3** *Let $B[X]$ be a restriction of $B$ to a trap space $X \in \mathcal{T}_{all}(B)$. Then the attractors $\mathcal{A}(B[X])$ extended with fixed values from $X$ are exactly the attractors of $B$ that intersect $X$:*

$$\{A_1 \in \mathcal{A}(B) \mid A_1 \subseteq \mathcal{S}(X)\} = \{A_2 \uparrow X \mid A_2 \in \mathcal{A}(B[X])\} \tag{8}$$

**Proof** Similar to Corollary S2.1, this easily follows from Corollary S1.1 and the fact that $X$ is a trap space. Since $STG(B[X])$ is isomorphic to the subgraph of $STG(B)$ induced by $\mathcal{S}(X)$ and there are no outgoing edges from $\mathcal{S}(X)$, every attractor within $\mathcal{S}(X)$ is also an attractor of $STG(B[X])$. $\square$

This lemma demonstrates that for a specific SD node $X$, we can further simplify the attractor detection problem by only focusing on the network $B[X]$.

#### S3.2 NFVS attractor candidate identification

Next, we describe how the attractor detection mechanism operates for a given $X \in V(SD(B))$. As stated previously, this part is based on the method from [22, 33], but proposes novel modifications that improve its scalability.

First, we need to define the notion of a *retained set*, and a Boolean network bound by a retained set. Formally, a retained set is any $R \in \mathbb{B}^n_\star$ (i.e., it is isomorphic to a subspace). However, its function is not to describe a set of states, but rather to be used as a limitation on the update functions of a Boolean network. We say that for a Boolean network $B$ and a retained set $R$, a network $B$ bound by $R$, written $B_{\downarrow R}$ is a network where variables fixed by $R$ can only change towards the value prescribed by $R$, but otherwise behave as originally defined by $B$. Formally, for any $v \in Var_n$ s.t. $R_v \in \mathbb{B}$, the update functions of network $B_{\downarrow R}$ are defined as follows:

$$f'_v(x) = (x_v \Rightarrow 1) \land (\neg x_v \Rightarrow f_v(x)) \qquad \text{if } R_v = 1$$
$$f'_v(x) = (\neg x_v \Rightarrow 0) \land (x_v \Rightarrow f_v(x)) \qquad \text{if } R_v = 0$$

Here, $f_v$ refers to the update function of $B$ and $f'_v$ to the update function of $B_{\downarrow R}$. Clearly, any variable $v$ fixed by $R$ cannot be updated by a cycle in $STG(B_{\downarrow R})$, because once it changes to the value $R_v$, it stays fixed forever. Furthermore, based on [22, 33], we know that if the variables fixed by $R$ represent a negative feedback vertex set of $IG(B)$, then the resulting $STG(B_{\downarrow R})$ only has fixed-point attractors, and that every attractor of $STG(B)$ is covered by at least one such fixed-point. Formally:

$$\forall A \in \mathcal{A}(B).\ \exists x \in \mathcal{T}_{fix}(B_{\downarrow R}).\ x \in A \tag{9}$$

This provides us with a set of *attractor candidate states*. By computing an NFVS $F$ of $IG(B[X])$ and choosing *some* retained set $R$ that fixes all variables prescribed by $F$, we obtain a network $B[X]_{\downarrow R}$ whose fixed-points cover all attractors of $B[X]$. Furthermore, for a fixed $X \in V(SD(B))$, we are only really interested in attractors that appear in $\mathcal{S}(X) \setminus \textsc{MotifStates}(X, B)$. Within network $B[X]$, we can thus safely ignore any fixed-points that appear in $\textsc{MotifStates}(\star^{n[X]}, B[X])$. These may still cover existing attractors, but these are attractors for which $\mathcal{T}(A) \neq X$.

Of course, there is no guarantee that these candidates correspond one-to-one with the relevant attractors, or even that each candidate covers *some* attractor (i.e., there can be spurious fixed-points that do not correspond to an attractor state). This will be addressed in subsequent steps of the method.

However, we should observe that when the candidate set is empty, it proves that there are no relevant attractors in $X \setminus \textsc{MotifStates}(X, B)$. Furthermore, if $X$ is a minimal trap space, then a singleton candidate set is also final, since each minimal trap space must contain at least one attractor. As such, if the candidate set is empty (or a singleton for a minimal trap space) the attractor seed state search is complete. In the following, we assume that the method checks for these conditions and we do not explicitly state them again.

#### S3.3 ASP-based retained set optimization

In the previous section, we observed that any retained set $R$ that fixes variables from an NFVS of $IG(B[X])$ is sufficient to produce candidate states that *cover* all attractors of $B[X]$. However, the choice of a particular $R$ can impact the size of the candidate set significantly. We are thus motivated to explore various options for selecting $R$ that lead to a minimal set of candidates (often empty).

In [22, 33], the authors propose a heuristic based on analysing the update functions of the Boolean network to obtain a good retained set. While this technique is reasonably effective, it can still result in candidate sets that are too large to enumerate.

To complement this method, we propose a different approach (Algorithm S4), based on incrementally building the retained set while monitoring the size of the resulting candidate set. Whenever the size of the candidate set grows, we greedily optimize the retained set and continue with a locally minimal result.

**def** BuildRetainedSet($X$ : target subspace)**:**
  $(R, c) \leftarrow (\star^{n[X]}, 0)$;
  **for** $v \in$ NFVS$(IG(B[X]))$ **do**
    $(R_0, R_1) \leftarrow (R[v \leftarrow 0], R[v \leftarrow 1])$;
    `/* Enumerations can interleave and terminate when the first one (i.e. smaller result) completes. */`
    $c_0 \leftarrow |\mathcal{T}_{fix}(B[X]_{\downarrow R_0}) \setminus \text{MotifStates}(\star^{n[X]}, B[X])|$;
    $c_1 \leftarrow |\mathcal{T}_{fix}(B[X]_{\downarrow R_1}) \setminus \text{MotifStates}(\star^{n[X]}, B[X])|$;
    $(R', c') \leftarrow$ **if** $c_0 \leq c_1$ **then** $(R_0, c_0)$ **else** $(R_1, c_1)$;
    **if** $c' > c$ **and** $c' \geq$ OptLimit **then**
      $(R', c') \leftarrow$ Optimize$(X, R')$;
    $(R, c) \leftarrow (R', c')$;
  **return** $R$;
**def** Optimize($X$ : target subspace, $R$ : candidate retained set)**:**
  $done \leftarrow false$;
  $c \leftarrow |\mathcal{T}_{fix}(B[X]_{\downarrow R}) \setminus \text{MotifStates}(\star^{n[X]}, B[X])|$;
  **while** $\neg done$ **do**
    $done \leftarrow true$;
    **for** $v \in Var_n$ s.t. $R_v \in \mathbb{B}$ **do**
      $R' \leftarrow R[v \leftarrow \neg R_v]$;
      `/* Enumeration can terminate early if` $c' \geq c$. `*/`
      $c' \leftarrow |\mathcal{T}_{fix}(B[X]_{\downarrow R'}) \setminus \text{MotifStates}(\star^{n[X]}, B[X])|$;
      **if** $c' < c$ **then**
        $done \leftarrow false$;
        $(R, c) \leftarrow (R', c')$;
  **return** $(R, c)$;

**Algorithm S4:** Algorithm for building the optimized retained set $R$ using iterative queries to the fixed-point problem.

In Algorithm S4, operation $|\mathcal{T}_{fix}(B[X]_{\downarrow R}) \setminus \text{MotifStates}(\star^{n[X]}, B[X])|$ is implemented by encoding the fixed-point problem as an ASP program, similar to what was presented in [31]), and enumerating the solutions using `clingo`. Excluding all the MotifStates$(\star^{n[X]}, B[X])$ solutions can be also done at the level of the ASP program (the program specifies that the solutions must not belong to any of the maximal subspaces of the universal space $\star^{m[X]}$). Note that since the problem is encoded as an ASP program and we only care about the number of solutions (i.e., stable models) of this program, we could also use some counting methods in ASP [34] (i.e., only count the number of stable models without enumerating them).

To compute the NFVS, we use a heuristic routine based on greedily covering the shortest uncovered negative cycles, implemented in AEON [32], meaning the NFVS is not necessarily optimal for all networks. Finally, to avoid unnecessary optimization when the result is already reasonably simple, `biobalm` provides a configurable OptLimit constant, which allows to skip optimization if it is deemed unnecessary (the default value is 1000).

Clearly, the method is correct in the sense that once it terminates, $R$ is a valid retained set that fixes every variable from an NFVS of $IG(B[X])$. Unfortunately, its complexity is obviously much greater than that of the heuristics from [22, 33]. However, as we show later in the experiments, it is typically capable of avoiding the candidate set explosion that can occur when using the original heuristics. To avoid spending unnecessary time on optimization in cases that are already easy to solve, we thus first use the original heuristic method with a hard limit on the number of candidates (the OptLimit), and only if this limit is exceeded, we proceed to use the incremental construction proposed in Algorithm S4.

#### S3.4 Simulation-based candidate set reduction

Once the retained set is selected and the attractor candidate states are computed, we aim to reduce this set as much as possible (unless it is already empty, or a singleton in case of a minimal trap space, per previous discussion). In [22, 33], the authors use stochastic simulation to eliminate spurious candidates. Here, we adapt this approach, but with an enhanced simulation algorithm that allows us to prune the candidate set more efficiently.

The approach is described in Algorithm S5. Here, Bdd($S$) generally refers to a binary decision diagram that encodes the characteristic Boolean function of the given set of states $S \subseteq \mathbb{B}^n$. For such a decision diagram $S_{bdd}$, we then write $S_{bdd}(x)$ to denote that the characteristic function evaluates to 1 (i.e., $x$ belongs to the original set $S$).

Compared to [22, 33], we present two improvements: First, the step limit for the simulation is exponentially increasing, as long as the method is still capable of eliminating spurious candidate states. This means that compared to [22, 33], the simulation can run for much longer, assuming it is actually achieving progress. Second improvement is the use of binary decision diagrams to represent the update functions and the sets of target states to optimize the inclusion tests for large candidate sets.

We discuss the correctness of SimAny and SimMin separately. In SimAny, the method computes *step* successor states by updating each eligible network variable. If any of these successor states appears in the *avoid* set, the simulation terminates and the state is not retained. If the simulation completes without encountering a state from the *avoid* set, the final state is returned as a member of $C'$. Meanwhile, in SimMin, all states are simulated one step at a time, with $C'$ representing the successors of $C$ after one step.

```
def Simulate(X : target subspace, C : candidate states):
    s ← 2^10;
    loop
        C′ ← if X is min. then SimMin(X, C, s) else SimAny(X, C, s);
        if |C′| = |C| then break;
        C ← C′;
        s ← 2 · s;
    return C;
def SimAny(X : target subspace, C : candidate states, steps: limit):
    update_v ← Bdd(f_v) for all v ∈ free(X);
    avoid ← Bdd(C) ∨ Bdd(MotifStates(X, B));
    C′ ← ∅;
    for c ∈ C do
        avoid ← avoid ∧ ¬Bdd({c});
        keep ← true;
        for 0 . . . steps do
            for v ∈ free(X) in randomized order do
                c_v ← update_v(c);
            if avoid(c) then
                keep ← false;
                break;
        if keep then
            avoid ← avoid ∨ Bdd({c});
            C′ ← C′ ∪ {c};
    return C′;
def SimMin(X : target subspace, C : candidate states, steps: limit):
    update_v ← Bdd(f_v) for all v ∈ free(X);
    avoid ← Bdd(C);
    for 0 . . . steps do
        C′ ← ∅;
        for c ∈ C do
            avoid ← avoid ∧ ¬Bdd({c});
            for v ∈ free(X) in randomized order do
                c_v ← update_v(c);
            if avoid(c) then continue;
            avoid ← avoid ∨ Bdd({c});
            C′ ← C′ ∪ {c};
        C ← C′
    return C;
```

**Algorithm S5:** Outline of the proposed simulation method. The actual simulation approach differs between minimal and non-minimal trap spaces, since the set MotifStates$(X, B)$ is empty in the former.

Note that in both methods, the new candidate set consist of the final simulated states, not the original input candidates. This ensures that the subsequent simulation run (if any) resumes with candidates in which the previous run ended. Furthermore, a state can be eliminated as a candidate only if we can show that it reaches one of the other candidate states, or one of the inner motif states (for non-minimal trap spaces). This ensures that every relevant network attractor is still covered by the candidate set.

#### S3.5 Static analysis candidate set reduction

Finally, [22, 33] uses the tool `pint` [35] to further reduce the candidate set by studying the reachability between candidates through static analysis. In `biobalm`, we also integrate `pint` as an optional attractor candidate processing step. In our experience, the previously described improvements typically reduce the candidate set to the extent where using `pint` is no longer necessary. We use `pint` in our testing to demonstrate this fact, but within `biobalm`, it is an optional dependency that is only used when requested (and installed). In pseudocode, we denote this step as procedure $\textsc{PintOpt}(X : \text{target subspace}, C : \text{candidate states})$.

#### S3.6 Symbolic identification of attractor sets

If the previously described methods cannot exactly identify attractor seed states for the succession diagram node $X$, `biobalm` employs a symbolic reachability method based on *saturation* [36] to fully identify the attractor set and thus conclusively reduce the attractor candidate states to the seed states which one-to-one correspond to the network attractors.

Saturation is a symbolic reachability technique for asynchronous systems where the set of reachable states is always expanded by updating one system variable at a time. Furthermore, saturation always updates variables in the order in which they appear in the symbolic encoding, which tends to minimize the size of the intermediate set representations.

As an optional by-product of this process, the full attractor set can be obtained. If desired, `biobalm` can then use this method to compute the attractor set for every discovered attractor in the succession diagram. However, this often increases the runtime significantly if the attractors are complex and do not have a compact symbolic BDD representation.

To further accelerate the saturation process for each candidate state, we prioritise the saturation of variables for which the candidate differs from some of the other candidates or the inner motif states. This ensures that we first update variables that *necessarily need to change*, assuming the candidate is to be eliminated.

The method is described in Algorithm S6. Here, we use the same $\textsc{Bdd}$ notation as in Algorithm S5, and further write $\textsc{Post}_v(S, B)$ to denote the BDD which represents the *successors* of set $S$ w.r.t. $STG(B)$ obtained by updating

```
def SymbolicSeedStates(X : subspace, C : candidate states):
    S_X ← ∅;
    for c ∈ C do
        C ← C \ {c};
        attr_bdd ← AttractorSet(c, X, C ∪ S);
        if attr_bdd ≢ 0 then
            /* If desired, attr can be also stored for further
               analysis, as it is exactly the attractor set
               reachable from the candidate state c.          */
            S_X ← S_X ∪ {c};
    return S_X;
def AttractorSet(x : candidate, X : subspace, C : other candidates):
    avoid ← Bdd(C) ∨ Bdd(MotifStates(X, B));
    attr ← Bdd({x});
    conflict ← { v ∈ free(X) | ∃y : avoid(y) ∧ x_v ≠ y_v };
    other ← free(X) \ conflict;
    /* Place conflict variables before other variables.    */
    remaining ← concat(sorted(conflict), sorted(other));
    satruate ← ∅;
    loop
        /* Saturate already selected variables.             */
        done ← false;
        while ¬done do
            done ← true;
            if (avoid ∧ attr) ≢ 0 then
                return 0;
            for v ∈ sorted(saturate) do
                step ← post_v(attr, B) ∧ ¬attr;
                if step ≢ 0 then
                    attr ← attr ∨ step;
                    done ← false;
                    break;
        /* Pick additional variable for the saturation loop,
           or terminate if all variables are saturated.     */
        if remaining is empty then
            return attr;
        for v ∈ remaining do
            if (post_v(attr, B) ∧ ¬attr) ≢ 0 then
                saturate ← saturate ∪ {v};
                remove(remaining, v);
```

**Algorithm S6:** The symbolic reachability method used for exact identification of attractor sets.

of variable $v$. See [37] for a more in-depth description of how such function is computed using BDDs.

Furthermore, in order for saturation to work optimally, it is necessary to update variables in the order in which they appear in the symbolic encoding. In Algorithm S6, this is implemented using the function SORTED. Finally, we assume that the variable *remaining* is an ordered list, not a set. Hence, when exploring this list, we first visit the $conflict$ variables, and then the *other* variables. This ensures that whenever possible, we first saturate the $conflict$ variables that are necessary to reach one of the *avoid* states. Note that in Algorithm S6, the reachability procedure only ever updates variables from $free(X)$, as the remaining variables are guaranteed to be fixed (due to $X$ being a trap space).

#### S3.7 Summary

To conclude, we present a summary of the attractor detection process (one can also refer to the corresponding figure of the main paper). The correctness of the whole method then hinges on the following observations:

1. Considering a (fully expanded) succession diagram $SD(B)$ and an attractor $A \in \mathcal{A}(B)$, there is exactly one $X \in V(SD(B))$ such that $A \subseteq (\mathcal{S}(X) \setminus \textsc{MotifStates}(X, B))$ (Lemma S3.2). As such, each subspace $X \in V(SD(B))$ can be considered independently when searching for attractors and states from $\textsc{MotifStates}(X, B)$ can be safely ignored. Furthermore, network $B[X]$ can be used to identify the attractors, instead of relying on the full network $B$ (Lemma S3.3).

2. Given a *retained set* $R$ which fixes all variables of some NFVS $F$ of $IG(B)$, the set of fixed-points $\mathcal{T}_{fix}(B_{\downarrow R})$ covers all attractors of $B$ (each original attractor contains at least one such fixed-point) [22, 33]. Following (1), this can be individually applied to each subspace $X \in V(SD(B))$ and the subspace-restricted network $B[X]$ (Section S3.2).

3. The choice of a particular retained set $R$ is arbitrary as long as it fixes all variables of some NFVS of $IG(B)$. As such, we can optimize the retained set to reduce the initial number of attractor candidate states $C = \mathcal{T}_{fix}(B[X]_{\downarrow R})$ (Section S3.3).

4. An attractor candidate state $x \in C$ is unnecessary for a subspace $X \in V(SD(B))$ if it (a) can reach any other subspace $Y \in V(SD(B))$ s.t. $Y \neq X$ (it cannot be a member of an attractor); or (b) can reach any other candidate state $x' \in C$ s.t. $x' \neq x$ ($x$ and $x'$ would appear in the same attractor, hence it is sufficient to retain $x'$). These conditions can be incompletely checked using simulation (Section S3.4) or static analysis (Section S3.5), further reducing the set of candidate state $C$.

5. If the candidate set for $X \in V(SD(B))$ is inconclusive (more than one candidate in a minimal trap spaces, or any candidate in a non-minimal

trap space), symbolic reachability can be used to exactly identify the smallest trap set containing each candidate $c \in C$. This enables us to test the rules of (4) exhaustively and eliminate any spurious or duplicate candidate states, obtaining the exact attractor seed states $\mathcal{A}_{seed}(B) = \bigcup_{X \in V(SD(B))} S_X$. Here, $S_X$ is the set of seed states identified for the subspace $X$ (Section S3.6).

6. Optionally, the same reachability procedure can be used to fully identify each attractor set in $\mathcal{A}(B)$ based on the seed states $\mathcal{A}_{seed}(B)$.

Overall, this leads us to the following conclusion:

**Theorem S3.1** *Let $B$ be a BN with a succession diagram $SD(B)$, let $A \in \mathcal{A}(B)$ be its attractor, and let $S_X$ denote the set of attractor seed states identified by the workflow of Section S3.7 for node (i.e. subspace) $X$ of $SD(B)$.*

1. *$\mathcal{T}(A) \in V(SD(B))$ for some subspace $X_A = \mathcal{T}(A)$.*
2. *$S_Y \cap A = \emptyset$ for every other $Y \neq X_A$.*
3. *$S_{X_A} \cap A = \{x_A\}$, where $x_A$ is the seed state of attractor $A$.*

**Proof** Intuitively, we claim that for each attractor $A$ and the smallest trap space $\mathcal{T}(A)$ that contains $A$, out method identifies within this trap space exactly one seed state $x_A$ from $A$ (the exact choice of $x_A$ is arbitrary). Claims one and two simply follow from Lemma S3.2 and the fact that by construction, $S_X \subseteq \mathcal{S}(X)$. The third claim is then argued in the discussion above: For every attractor that appears in $X \setminus \textsc{MotifStates}(X, B)$ ($A$ being one of them), we first identify a set of candidate states $C_X$ for which we have $A \cap C_X \neq \emptyset$. Subsequently, we reduce this $C_X$ by various methods until $|A \cap C_X| = 1$. □

### S4 Control strategy identification

Previous work [38, 11, 9] has produced several algorithms that use the succession diagram for target attractor control. These algorithms share a common approach: a path in the succession diagram from the root node to a target trap space is identified, and a control strategy for each node along the identified path is selected. The various algorithms differ primarily in how the control strategy for each succession diagram node is obtained, but a simple brute-force approach is usually sufficient. In `biobalm`, we implement the internal history and minimal history algorithms described in [11] with only slight modifications to allow for dynamic expansion of the succession diagram. Specifically, if a succession diagram node is not expanded, the control algorithms will only expand it if it contains the target trap space. This allows for partially solving the attractor control problem in networks with very large succession diagrams, provided the target attractor is known.

### S5 Partial SD expansion for attractor detection

While the method presented in Section S3 is complete, it always considers a fully expanded succession diagram. This could be considered wasteful, as most attractors typically appear within minimal trap spaces. As such, we may want to prioritize expanding the succession diagram only up to the point where all minimal trap spaces are discovered, assuming we can prove for the remaining (non-minimal) trap spaces that do not contain any further attractors.

In other words, we want to identify a succession diagram expansion method that guarantees the conclusions outlined in Theorem S3.1 without always fully expanding the succession diagram. In the following, we describe a method that achieves this by prioritising certain structurally independent parts of the Boolean network during the expansion process. However, it should be noted that this is by no means the only possible approach. Deeper knowledge of succession diagrams could inspire further investigation into which trap spaces are in fact relevant for attractor detection.

In the following, we use $\cap$ to denote the trap space that is the intersection of two trap spaces. In all instances, we either explicitly argue that the intersection is not empty, or this should be clear from context.

**Definition S5.1** *Let $V \subseteq Var_n$ be a set of network variables and $IG(B)$ an influence graph of a Boolean network. Then, we write $\mathcal{D}(V)$ to denote the subset of network variables upon which the set $V$ depends, including transitive dependencies. We call such $\mathcal{D}(V)$ the* source block *of $V$.*

Intuitively, we call $\mathcal{D}(V)$ the source block because it defines a sub-network of $B$ that does not depend on anything else in $B$, but can influence other network components (similar to a more commonly used term *source node*). Note that for a specific $V$, $\mathcal{D}(V)$ can in fact correspond to a single source node or a single source component of the influence graph. However, it can also be a larger collection of network nodes.

**Example S5.1** *Recall the Boolean network from our running example (Example S1.1) and its influence graph, shown in Figure S2. The source block of $\{v_4\}$ is the set $\mathcal{D}(\{v_4\}) = \{v_1, v_2, v_4\}$. Meanwhile, the source block of $\{v_1, v_2\}$ is $\mathcal{D}(\{v_1, v_2\}) = \{v_1, v_2\}$, because these states already represent a source SCC of the influence graph.*

#### S5.1 Source blocks and network restrictions

First, we observe an important property that allows us to relate the behaviour of the source block variables with the rest of the network:

**Lemma S5.1** *Let $C$ be a source block of $B$, and let $X_1$, $X_2$ be any two subspaces such that $C = free(X_1) = free(X_2)$. Then the sub-graph of $STG(B)$ induced by $\mathcal{S}(X_1)$ is isomorphic to the sub-graph induced by $\mathcal{S}(X_2)$.*

Conversely, this lemma states that by fixing all variables other than the ones in $C$, we obtain the same behaviour (same STG sub-graph), regardless of the values to which we fix those variables. To prove it, assume that the isomorphism maps every state from $\mathcal{S}(X_1)$ to the state in $\mathcal{S}(X_2)$ with the same values of free variables in $C$. Formally, we can define this as $id(x) = (x \downarrow X_1) \uparrow X_2$ (for any $x \in \mathcal{S}(X_1)$).

Now, consider an STG transition $x_1 \to x_2$ such that, w.l.o.g, this transition appears in the sub-graph induced by $\mathcal{S}(X_1)$, but not in the sub-graph of $\mathcal{S}(X_2)$. First, this transition must update some $v \in C$, since all other variables are fixed. However, the update function of such $v \in C$ only depends on other variables $v' \in C$ (i.e. $in(f_v) \subseteq C$). Thus the output of $f_v$ is the same also for the corresponding state $x_1' = id(x_1)$ within $\mathcal{S}(X_2)$, as this state has the same configuration of the free variables of $C$, i.e. $f_v(x_1) = f_v(id(x_1))$. $\square$

This property allows us to define $B[C]$, i.e. network $B$ restricted to the source block $C$, by which we mean $B[C] = B[C^0]$, where $C^0$ is a subspace s.t. $free(C^0) = C$ and every other variable is set to 0. Such network $B[C]$ then describes the behaviour of variables within $C$, while ignoring the rest of the network that has no influence on $C$.

Note that similar notation could be in theory considered also for subsets of $Var_n$ that are not source blocks. However, for source blocks, we have just shown that the exact choice of the fixed values is in practice irrelevant, since it produces the same STG sub-graph (hence we default to the subspace $C^0$). For other sets, different choice of fixed values can lead to different behaviour.

#### S5.2 Source blocks and trap spaces

First, we observe a helpful lemma which hints at the relationship between source blocks and maximal trap spaces of a Boolean network:

**Lemma S5.2** *Given any set $V \subseteq Var_n$ and any (globally) maximal trap space $X \in \mathcal{T}_{max}(B)$, it holds that either $fixed(X) \subseteq \mathcal{D}(V)$, or $fixed(X) \cap \mathcal{D}(V) = \emptyset$.*

**Proof** For a contradiction, assume this does not hold. That is, $fixed(X) \cap \mathcal{D}(V) = V'$ such that $V' \neq \emptyset$ and $V' \neq fixed(X)$. Then, let $X'$ be a space such that $X'_v = X_v$ for all $v \in V'$ and $X'_v = \star$ otherwise (i.e. all fixed variables that are not within $\mathcal{D}(V)$ are set to $\star$). In the next paragraph, we argue that such $X'$ is in fact a trap space, which completes the proof by contradiction, as it violates the claim that $X$ is maximal, since $X'$ is clearly a super-space of $X$.

To show that $X'$ is indeed a trap space, let us consider a state $x \in \mathcal{S}(X')$ and a variable $v \in Var_n$ such that $x \to_v y$ (i.e. $x$ can transition into state $y$ by updating variable $v$). First, if $x \in \mathcal{S}(X)$, we know that $y \in \mathcal{S}(X)$ and by extension $y \in \mathcal{S}(X')$, since $X$ is a trap space and $X'$ is its superset. Therefore, assume $x \in \mathcal{S}(X') \setminus \mathcal{S}(X)$. Now, if $v \in free(X')$, then $y \in \mathcal{S}(X')$, because $X'_v = \star$, so we cannot escape $X'$ by updating a free variable. Finally, assume that $v \in fixed(X')$, which would imply that $y \notin \mathcal{S}(X')$. Then, we know that the update function $f_v$ only depends on variables from $\mathcal{D}(V)$, i.e. $in(f_v) \subseteq \mathcal{D}(V)$.

Since $fixed(X') \cap \mathcal{D}(V) = fixed(X) \cap \mathcal{D}(V)$, we can modify the state $x$ by fixing all variables that do not belong to $\mathcal{D}(V)$ in accordance with $X$ without modifying the output of $f_v$. In other words, for any such $x \in \mathcal{S}(X')$, there exists an $x' = (x \downarrow X') \uparrow X \in \mathcal{S}(X)$ that agrees with $x$ on all variables within $\mathcal{D}(V)$ and hence agrees on the output of $f_v$, $f_v(x) = f_v(x')$. Finally, notice that $v \in fixed(X)$ (and $v \in \mathcal{D}(V)$, thus $x_v = x'_v$), which contradicts the assumption that $X$ is a trap space. □

Intuitively, what this lemma states is that each maximal trap space $X$ can be associated with a set of source blocks that contain all fixed variables of $X$ and there cannot be any overlap between the fixed variables of $X$ and other source blocks (each source block either covers $X$ fully, or not at all).

To further refine this relationship, we observe that $\mathcal{D}(fixed(X))$ is clearly the smallest source block (in terms of set inclusion) that covers the whole set $fixed(X)$. We call $\mathcal{D}(fixed(X))$ the *canonical source block* of $X$, written $\mathcal{CD}(X)$. Such source block can also cover other maximal trap spaces $X'$, in which case (by Lemma S5.2), we get that $\mathcal{CD}(X') \subseteq \mathcal{CD}(X)$. Note that it is still possible for some combinations of $X$ and $X'$ that $\mathcal{CD}(X') \subset \mathcal{CD}(X)$. However, this implies that $fixed(X') \cap fixed(X) = \emptyset$.

Before we move further, we also state the following property, which formalizes the fact that trap spaces from one source block can be combined with any other trap space that the source block does not depend on:

**Lemma S5.3** *Let $X$ and $Y$ be maximal trap spaces s.t. $\mathcal{CD}(X) \cap fixed(Y) = \emptyset$. Then $Z = X \cap Y$ is also a trap space (albeit not maximal).*

**Proof** This trivially follows from the fact that $\mathcal{CD}(X) \cap fixed(Y) = \emptyset$ also implies $fixed(X) \cap fixed(Y) = \emptyset$, which means that $Z = X \cap Y$ is well defined. Subsequently, any intersection of two trap spaces is also a trap space. □

Now, we can establish several useful properties of the canonical source blocks:

**Proposition S5.1** *Let $C \subseteq Var_n$ be a canonical source block of some $X_C \in \mathcal{T}_{max}(B)$ and let $X_1, \ldots, X_k \in \mathcal{T}_{max}(B)$ be all the maximal trap spaces for which $fixed(X_i) \subseteq C$. Then, for every minimal trap space $Y \in \mathcal{T}_{min}(B)$, there exists an $X_i$ s.t. $\mathcal{S}(Y) \subseteq \mathcal{S}(X_i)$ ($Y$ is a subspace of $X_i$).*

**Proof** First, note that (as per the discussion above), it does not necessarily hold that for each $X_i$, we have $\mathcal{CD}(X_i) = C$. We only know that $\mathcal{CD}(X_i) \subseteq C$. Second, note that the proposition does not explicitly cover the trivial case of $\mathcal{T}_{max} = \emptyset$, as there is only one minimal trap space $\star^n$. Such case is therefore generally not interesting as the full succession diagram has a single node.

Now, for a contradiction, let there be some $Y \in \mathcal{T}_{min}(B)$ such that for every $X_i$, $X_i \cap Y = \emptyset$. Furthermore, we assume that $fixed(Y) \cap C \neq \emptyset$, otherwise by Lemma S5.3, $Y$ is not minimal as we can intersect it with any $X_i$, yielding a smaller trap space.

Next, we argue that a space $Y'$ which fixes all variables in $C$ according to $Y$ is a trap space (i.e. $Y'_v = Y_v$ if $v \in C$ and $Y'_v = \star$ otherwise). This follows (similar to the proof of Lemma S5.2) from the fact that variables within $C$ only depend on other variables in $C$. Hence if $Y'$ were not to be a trap space, there must be a state disproving this and we can modify this state by fixing the variables in $fixed(Y) \setminus C$ to match the requirements of $Y$ (since these variables do not matter to the update functions of any $v \in C$, we can safely fix them to arbitrary values). Finally, $Y'$ is clearly a super-space of $Y$ and $fixed(Y') \subseteq C$. Thus, by the nature of set inclusion and Lemma S5.2, either $Y'$ is a sub-space of some $X_i$, or it is itself a maximal trap space and $Y' = X_i$ for some $i \in [1, k]$. Here, Lemma S5.2 is used to argue that there cannot be any other maximal trap space $Z$ s.t. $Y' \subseteq Z$, $fixed(Z) \cap C \neq \emptyset$, yet $Z \notin X_1, \ldots X_k$. Consequently, $Y$ has a super-space within the spaces $X_1, \ldots, X_k$. □

Intuitively, we have just shown that if we take any canonical source block $C$, then the trap spaces which fix variables from this block are sufficient to reach any minimal trap space within the succession diagram. In the context of a partially expanded succession diagram, it is thus always sufficient to only expand the successor nodes corresponding to one of the canonical source blocks in order to eventually expand all minimal trap spaces.

For simplicity, this section only talked about *globally* maximal trap spaces and their corresponding source blocks. However, the same conclusions can be applied to trap spaces that are *locally* maximal within a larger trap space $X$ (i.e. the set $\mathcal{T}_{max}(B, X)$). In particular, it is clear that all of the properties already hold in the network $B[X]$.

**Corollary S5.1** *Lemma S5.2, Lemma S5.3 and Proposition S5.1 also generalize to $\mathcal{T}_{max}(X, B)$ (as opposed to just $\mathcal{T}_{max}(B)$) by the means of network $B[X]$.*

We simply need to consider that $\mathcal{T}_{max}(X, B)$ one-to-one correspond to the $\mathcal{T}_{max}(B[X])$ of a restricted network $B[X]$ (Corollary S2.1). In the corresponding propositions, we then also consider the influence graph $IG(B[X])$ instead of the full $IG(B)$ to compute which variables belong to the relevant source blocks.

#### S5.3 Source blocks and attractors

Next, we explore the relationship between source blocks and attractors of a Boolean network. We already know that each minimal trap space contains at least one attractor and thus needs to be explored. However, other trap spaces may or may not contain additional attractors. Here, we formulate a condition which allows us to simplify the attractor search in such non-minimal trap spaces.

**Proposition S5.2** *Let $C \subseteq Var_n$ be a canonical source block of some $X_C \in \mathcal{T}_{max}(B)$ and let $S_X$ denote the attractor seed states computed for SD node $X$.*

*Let $SD(B)$ be the succession diagram of $B$ and $R$ its root node. Furthermore, let $SD(B[C])$ be the succession diagram of the network $B$ restricted to $C$, and let $R_C$ be its root node (i.e. $R_C = R \downarrow C^0$)).*

*If $S_{R_C} = \emptyset$ (root node of $SD(B[C])$ has no attractor seeds), then for every $A \in \mathcal{A}(B)$, it holds that $A \subseteq X_A$ for some $X_A \in \mathcal{T}_{max}(B)$ s.t. $\mathcal{CD}(X_A) \subseteq C$. In such case, we also have that $S_R = \emptyset$ (root of $SD(B)$ has no attractor seeds).*

**Proof** Intuitively, this proposition claims that if the root of $SD(B[C])$ contains no attractors, then the root of $SD(B)$ also contains no attractors, and furthermore, all attractors of $B$ appear in maximal trap spaces whose canonical source blocks are subsets of $C$. Importantly, this holds for any $C$ that is a canonical source block of some maximal trap space.

To prove this, first, let us first recall that by definition, the update functions of variables in $C$ are the same in $B$ and $B[C]$, since they only depend on variables from $C$ (i.e. eliminating the other variables does not impact them at all). Then, let $X_1, \ldots, X_k \in \mathcal{T}_{max}(B)$ be all the maximal trap spaces for which $fixed(X_i) \cap C \neq \emptyset$ (and thus $\mathcal{CD}(X_i) \subseteq C$). We claim that $\mathcal{T}_{max}(B[C]) = \{X_1, \ldots, X_k\} \downarrow C^0$. That is, the maximal trap spaces of the restriction $B[C]$ are exactly the projections of maximal trap spaces whose canonical source block is a subset of $C$. This easily follows from Lemma S5.2 and the definition of a canonical source block: For each maximal trap space $X \in \mathcal{T}_{max}(B)$, either (a) $\mathcal{CD}(X) \cap C = \emptyset$, in which case none of the $fixed(X)$ variables are present in $B[C]$; or (b) $\mathcal{CD}(X) \subseteq C$, in which case all variables that appear in $fixed(X)$ appear in $B[C]$ unchanged. Also, (c), no new larger trap spaces can appear in $B[C]$ as these would have to also be present in $B$, since their update functions would be the same.

Next, we observe that if the root $R_C$ of $SD(B[C])$ does not contain any proper attractors ($S_{R_C} = \emptyset$), it means that for every (restricted) network state $x \in (\mathcal{S}(R_C) \setminus \textsc{MotifStates}(R_C, B[C]))$, there exists a path $x \to y_1 \to \ldots \to y_m$ such that $y_m \in \mathcal{S}(X_i \downarrow C^0)$ for some $i \in [1, k]$ (i.e. $x$ eventually reaches a state in one of the projected maximal sub-spaces). However, the same path also exists for any state $x \in \mathcal{S}(R) \setminus \bigcup_{i \in [1,k]} \mathcal{S}(X_i)$ in the "full" succession diagram $SD(B)$, as we can extend every state $y_i$ along this path with *an arbitrary* fixed valuation of variables from $Var_n \setminus C$. Such path is clearly still valid, as none of the transitions depend on any variable in $Var_n \setminus C$.

To summarise, maximal trap spaces of $B[C]$ are exactly the projections of maximal trap spaces of $B$ whose fixed variables only depend on $C$. As such, if $S_R = \emptyset$, then every state in the root subspace either belongs to some of these $X_i$ (for $i \in [1, k]$), or it has a path into one such $X_i$. Consequently, every attractor $A \in \mathcal{A}(B)$ must appear in one of these maximal trap spaces $X_i$ whose fixed variables only depend on variables from $C$. $\square$

As before, we extend this proposition with the following corollary:

**Corollary S5.2** *Proposition S5.2 also generalizes to attractors in locally maximal trap spaces $\mathcal{T}_{max}(X, B)$ by evaluating it on the restricted network $B[X]$.*

As before, we simply need to consider that $\mathcal{T}_{max}(X, B)$ one-to-one correspond to the $\mathcal{T}_{max}(B[X])$ of $B[X]$ (Corollary S2.1) and that we should then consider the influence graph $IG(B[X])$ instead of the full $IG(B)$ to compute which variables belong to which relevant source blocks.

This proposition has two practical consequences for our attractor detection method: First, it allows us to rule out the presence of attractors in non-minimal trap spaces based on the attractors of network $B[C]$, which can be often much smaller than the full network $B$. Second, it complements Proposition S5.1 in that expanding only the maximal trap spaces of one source block $C$ not only eventually visits all minimal trap spaces, but (assuming $S_{R_C} = \emptyset$), all attractors are eventually found as well.

Finally, note that the presence of an attractor in the SD root node of subnetwork $B[C]$ does not *guarantee* that the corresponding trap space in the full succession diagram also contains an attractor. Updates of other variables within $Var_n \setminus C$ (and thus other canonical source blocks that do not intersect $C$) could be used to eventually reach one of the other maximal sub-spaces. In particular, since multiple canonical source blocks can exist (for different maximal trap spaces), it is sufficient if one of them rules out the existence of an attractor.

#### S5.4 Source block SD expansion

Finally, we can formulate how to apply the previously presented properties when designing a succession diagram expansion method. This method is presented as Algorithm S7.

```
def BlockExpand(X : SD node):
    if expanded(X) then return;
    Expand(X) // Algorithm S3
    if X is a minimal trap space then return;
    Y_1, ..., Y_m ← T_max(X, B);
    /* We could consider various criteria for selecting the
       canonical source block C.  Here, we choose the
       smallest C, since it also produces the smallest B[C].
       */
    C ← CD(Y_i) with minimal |CD(Y_i)|;
    B[C] ← restriction of B to variables from C;
    if AttractorSeeds(SD(B[C]), ⋆^|C|) = ∅ then
        /* By extension, no attractors can appear in X.     */
        AttractorSeeds(SD(B), X) ← ∅;
        for Y_i s.t. CD(Y_i) = C do
            BlockExpand(P^∞(Y_i));
    else
        /* Attractors in X not ruled-out by B[C].           */
        for Y_i ∈ Y_1, ..., Y_m do
            BlockExpand(P^∞(Y_i));
```

**Algorithm S7:** Partial SD expansion based on canonical source blocks.

First, let us observe some notable features of this algorithm: Algorithm S7 recursively traverses the succession diagram in a depth-first manner, as opposed to Algorithm S3 where we only described how to fully expand one SD node. After the algorithm terminates, the succession diagram is not necessarily fully expanded (Expand hasn't been called for all nodes). This is due to the first branch of the IF-condition, in which we intentionally skip the expansion of maximal sub-spaces where $\mathcal{CD}(Y_i) \neq C$ (by selecting the smallest $\mathcal{CD}(Y_i)$, we also ensure no other $\mathcal{CD}(Y_j)$ can be a subset of $C$).

Also observe that Algorithm S7 always terminates, because each iteration that recursively calls BlockExpand also expands one SD node. Since each SD is by definition finite, the recursion must eventually stop.

**Theorem S5.1** *For every attractor $A \in \mathcal{A}(B)$, the succession diagram generated with Algorithm S7 contains $X \in V(SD(X))$ s.t. $\mathcal{T}(A) = X$. Furthermore, this node $X$ is always an expanded node.*

**Proof** This follows from Proposition S5.1 and S5.2. First, the theorem is satisfied for any attractor that appears in a minimal trap space: Proposition S5.1 argues that it is always sufficient to expand the maximal trap spaces corresponding to one canonical source block to reach every minimal trap space. In Algorithm S7, both branches of the IF-condition expand (at least) the maximal trap spaces of the canonical source block $C$, so each minimal trap space is eventually found and expanded.

Second, any attractor $A$ for which $\mathcal{T}(A)$ is not minimal is covered by Proposition S5.2. In particular, if the root node of the restricted succession diagram $SD(B[C])$ does not have any attractor seeds, then we know that all attractors within $X$ reside in one of the maximal trap spaces corresponding to the source block $C$, and it is sufficient to expand those (and set the set of attractor seeds to $\emptyset$). In any other case, we expand *all* maximal trap spaces, since we cannot rule-out attractors in $X$ or any of its sub-spaces. □

#### S5.5 Expansion with size limit

In extreme cases, even the presented source block expansion can produce a succession diagram that is too large to enumerate. In that case, we allow for the expansion to be terminated early (based on a limit on the number of nodes). To facilitate attractor detection, we then connect each of the remaining nodes that have not been expanded yet directly to the minimal trap spaces contained in it (instead of maximal trap spaces). With this modification, it is still possible to detect every attractor. However, for a motif-avoidant attractor $A$, we can't guarantee that the succession diagram contains exactly the trap space $\mathcal{T}(A)$. As such, it could happen that more than one of the nodes where the expansion was terminated early (each now connected to its minimal trap spaces) contains the same motif-avoidant attractor. We use this method in our experiments to compute attractors in a small subset of very complex models. However, we did not find any motif-avoidant attractors, so our results are exact in this scenario.

#### S5.6 Summary

In this section, we have presented a method which is able to skip the expansion of certain succession diagram nodes while still guaranteeing that every minimal trap space and every attractor is properly accounted for. To do this, the method identifies independent "source blocks" within the influence graph $IG(B)$. Since these blocks of variables generate independent maximal trap spaces, it is sufficient to only expand one block at a time to eventually reach all minimal trap spaces. Furthermore, by analyzing the attractors of the sub-network corresponding to this block, we can typically rule out the presence of attractors in the main network. This means that it is generally sufficient to apply the attractor detection from Section S3 to the much smaller sub-networks, which provides further speed-up. For the trap spaces where the presence of attractors cannot be ruled out, the method falls back to the standard attractor detection process.

### S6 Benchmarks and experimental evaluation

In `biobalm`, we present a methodology that is generally applicable to a wide range of realistic Boolean networks and scalable beyond the capabilities of existing tools. As such, our testing considers two major datasets of networks.

#### S6.1 Biodivine Boolean Models

As described in [39], the Biodivine Boolean Models (BBM) dataset represents a comprehensive collection of realistic networks that are suitable for benchmarking and validation of Boolean network tools. At the time of writing, this covers the following sources:

- Biomodels [40] (Boolean and multi-valued binarized);
- CellCollective model repository [41] (Boolean);
- GINsim model repository [42] (Boolean and multi-valued binarized);
- COVID-19 disease map project [43] (Boolean);
- Kadelka dataset [44] and other independently source models (Boolean);

Collectively, this represents 230 Boolean networks ranging up to 342 variables and 1100 regulations. Since many networks contain constant input nodes that significantly influence the network's behavior (and therefore the structure of the succession diagram), for each model, we consider up to 128 variants with random (but distinct) constant input valuations. This process resulted in 14010 distinct benchmark instances.

#### S6.2 Random models

To test larger network sizes and networks with otherwise unusual properties, we also consider several ensembles of random networks.

**Critical N-K networks** We used `pystablemotifs` [11] to construct ensembles of random N-K models with $N \in \{10, 20, 40, 80, 160, 320, 640, 1280, 2560\}$ and $K \in \{2, 3\}$ (these values of $K$ are generally considered the most biologically realistic; for example, the BBM dataset has an average $K$ of $\sim 2.5$). The activation bias $p$ was set such that the sampled networks are in the critical regime (i.e. $p = 0.5$ for $K = 2$ and $p = 0.211325$ for $K = 3$). For each combination of values, we sampled 100 networks, giving 900 $K = 2$ and 900 $K = 3$ instances.

**Nested canalyzing networks** We also sampled random models with nested canalyzing functions (NCF) and $N \in \{10, 20, 40, 80, 160, 320, 640, 1280, 2560\}$, as these should better resemble real-world biomolecular networks. For these NCF models, we constructed the interaction graph with power-law out-degree and Poisson in-degree. We assigned 24% of edges to be negative, while having and 10% of the nodes as sink nodes. These choices align with the results of [44]. The average $K$ was set close to 2.5, which is the average $K$ of the BBM ensemble. The power of the out-degree distribution was varied with network size so that the distribution of $K$ is not correlated with the network size, as shown in [44]. We noticed that the NCF models had larger variance in $K$ compared to the critical N-K networks, which aligns with the observations for the BBM ensemble. This ensemble represents 900 unique benchmark instances.

**Dense random networks** Finally, we also consider 60 random networks originally used for the testing of `mts-nfvs` [8]. These are N-K networks with $K = 2$, but each function is chosen such that both of its inputs are essential. Consequently, the influence graph of these networks is typically strongly connected and generally does not have many symmetries or opportunities for reduction. The network sizes are $N \in \{100, 200, 300\}$, each cohort having 20 networks.

#### S6.3 Benchmark conditions

For any presented performance claims we consider a Ryzen 5800X CPU fixed to 4.7Ghz (with disabled frequency scaling) and 128GB of DDR4-3200 RAM.

Each experiment is using a single CPU core (none of the presented algorithms are parallelized), but we run up to four experiments in parallel to speed up the benchmark process. Each benchmark is limited to 32GBs of memory and an hour of runtime. Experiments were performed on a Debian 12 virtual machine using Python 12 and `clingo` 5.6 as the underlying ASP solver. The exact versions of the remaining dependencies are specified in the artefact.[3]

We evaluate three tools, `biobalm`, `AEON.py` [32] and `mts-nfvs` [8] on the attractor seed enumeration problem. For `biobalm`, we specifically consider the partial source block expansion instead of constructing the full succession diagram, and we cut-off the diagram construction at 1000 nodes. In each network, any constant values are first percolated to provide comparable initial conditions for each method.

[3] `https://doi.org/10.5281/zenodo.13854760`

We have not tested `pystablemotifs` in attractor identification, as previous results show that both `AEON.py` and `mts-nfvs` perform better. However, we have compared `pystablemotifs` to `biobalm` on full succession diagram construction and target control. Since the performance differential between the two tools is substantial, we only performed this experiment on the BBM models with input nodes fixed to *true* and a timeout of 10 minutes (we also excluded 4 models where succession diagram construction by either tool takes more than 10 minutes, yielding 226 benchmark instances).

##### S6.4 Benchmark results

In Figure S6, S7 and S8, we present two types of results for the attractor identification problem: First, the amount of benchmarks completed overall until a timeout. Second, the relative performance on these benchmarks, comparing `biobalm` to the competing tools.

Later, Figure S9 and Figure S10 show that `biobalm` clearly outperforms `pystablemotifs` at both succession diagram construction and control strategy identification.

#### S7 Results

This section presents the results of our structural analysis of large ensembles of biological and random succession diagrams.

##### S7.1 Networks with very large SD size

Table S1 lists the BBM networks that had more than 1000 nodes on average over the sampled input configurations in the fully expanded succession diagram. The listed number of nodes in the fully expanded succession diagram (sd size), depth of the succession diagram, and the number of minimal trap spaces are averaged over the sampled input configurations. All of them are curated.

##### S7.2 Attractor scaling

We enumerated all the attractors of the BBM networks, random critical N-K networks with K=2 and 3, and the NCF networks. Figure S11 shows that empirical models tend to be more heterogeneous than the random networks in terms of the number of attractors. The number of attractors is equal to the number of minimal trap spaces for each case, as there were no motif-avoidant attractors in any of the networks. The mean, variance, and kurtosis of the distribution is statistically significantly different between the BBM networks (averaged over input configurations) and the random networks, using bootstrap method with 95% confidence interval.

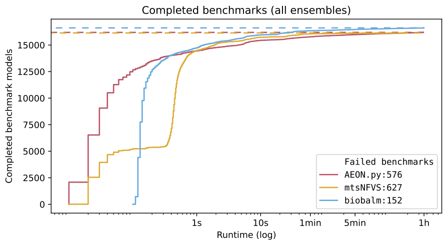


(a) All ensembles

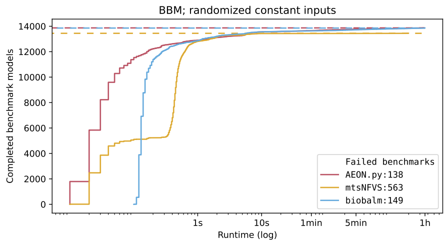


(b) Biodivine Boolean Models ensemble

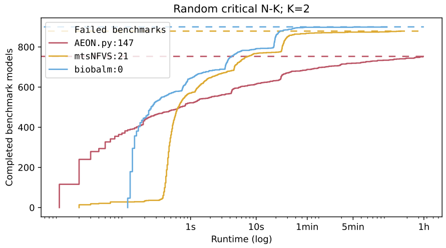


(c) Random critical N-K networks; $K = 2$

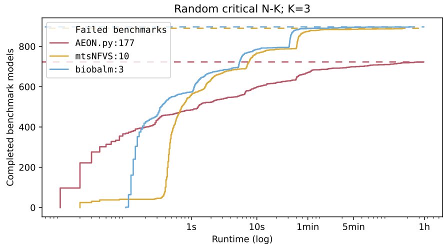


(d) Random critical N-K networks; $K = 3$

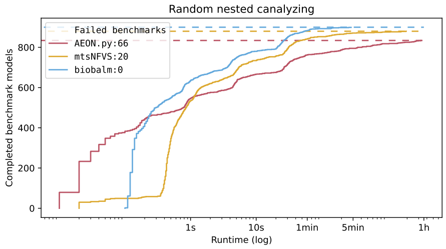


(e) Random nested-canalyzing networks

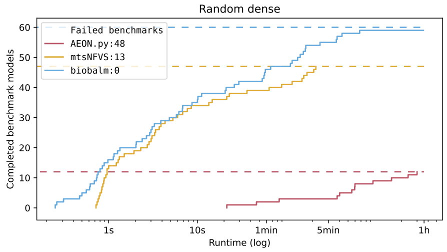


(f) Random dense networks ensemble

Figure S6: Cumulative completed benchmarks (until a certain time limit), comparing `biobalm` with `AEON.py` and `mts-nfvs`. Note that the time axis are logarithmic.

| BBM no. | name | nodes | sd size | depth | min. trap | citation |
|---|---|---|---|---|---|---|
| 002 | SIGNAL TRANSDUCTION IN FIBROBLASTS | 139 | 40125.8 | 16.7 | 303.1 | [45] |
| 143 | BREAST CANCER INHIBITORS | 97 | 34666.0 | 10.1 | 560.9 | [46] |
| 038 | SKBR3 BREAST CELL LINE LONG TERM | 25 | 17393.6 | 13.8 | 199.1 | [47] |
| 079 | COLORECTAL TUMORIGENESIS | 197 | 11560.9 | 11.6 | 46.7 | [48] |
| 034 | HCC1954 BREAST CELL LINE LONG TERM | 23 | 5212.9 | 13.6 | 147.7 | [47] |
| 033 | BT474 BREAST CELL LINE LONG TERM | 24 | 3544.0 | 10.7 | 52.3 | [47] |

Table S1: BBM networks that had more than 1000 nodes on average over the sampled input configurations in the fully expanded succession diagram. The listed number of nodes in the fully expanded succession diagram (sd size), depth of the succession diagram, and the number of minimal trap spaces (min. trap) are averaged over the sampled input configurations.

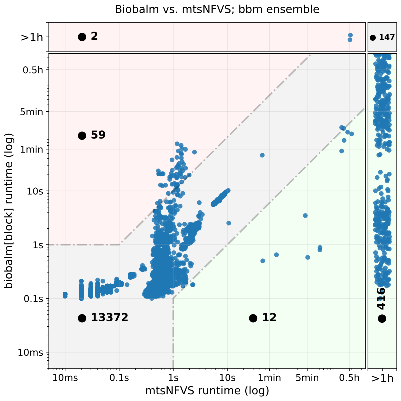


(a) `mts-nfvs`, BBM networks

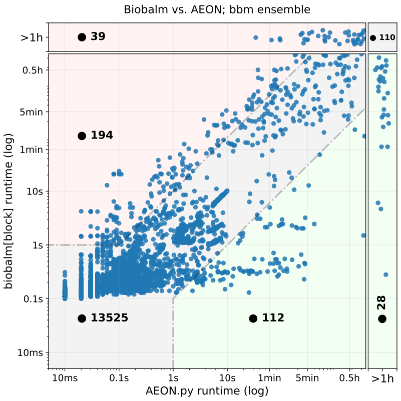


(b) `AEON.py`, BBM networks

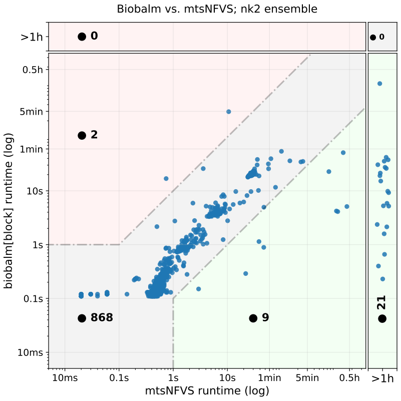


(c) `mts-nfvs`, critical $K = 2$

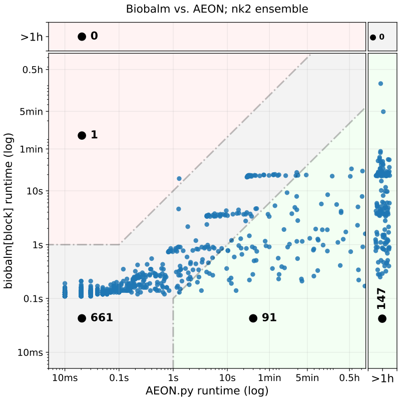


(d) `AEON.py`, critical $K = 2$

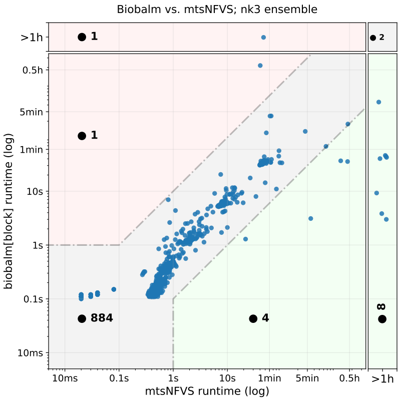


(e) `mts-nfvs`, critical $K = 3$

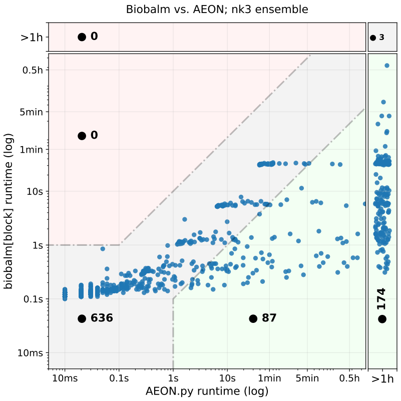


(f) `AEON.py`, critical $K = 3$

Figure S7: (Part one) Comparative performance of the considered tools vs. `biobalm` on the individual ensembles. Note that the time axis are logarithmic. The areas at the edges of the plot show models where the corresponding tool did not finish the benchmark.

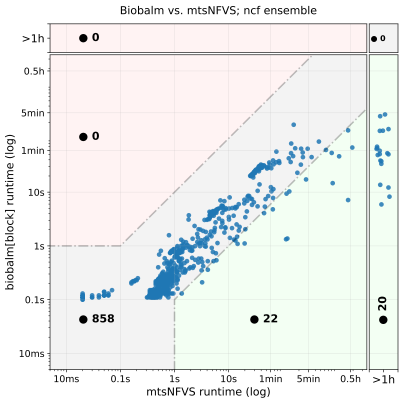


(a) **mts-nfvs**, nested canalyzing

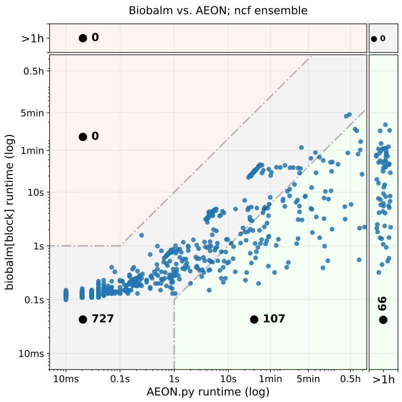


(b) **AEON.py**, nested canalyzing

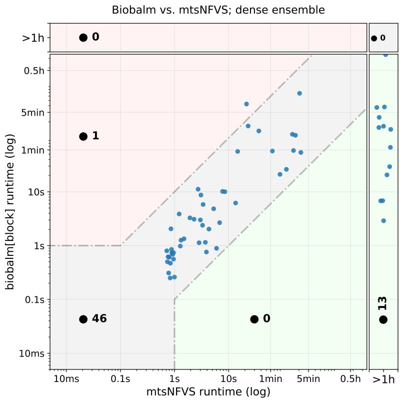


(c) **mts-nfvs**, dense

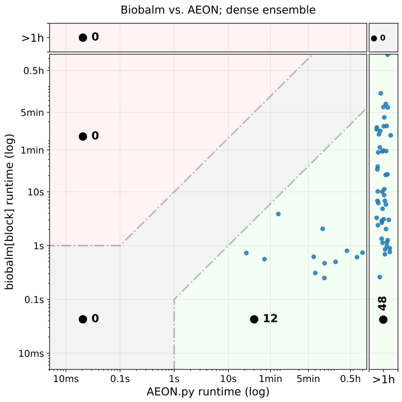


(d) **AEON.py**, dense

Figure S8: (Part two) Comparative performance of the considered tools vs. **biobalm** on the individual ensembles. Note that the time axis are logarithmic. The areas at the edges of the plot show models where the corresponding tool did not finish the benchmark.

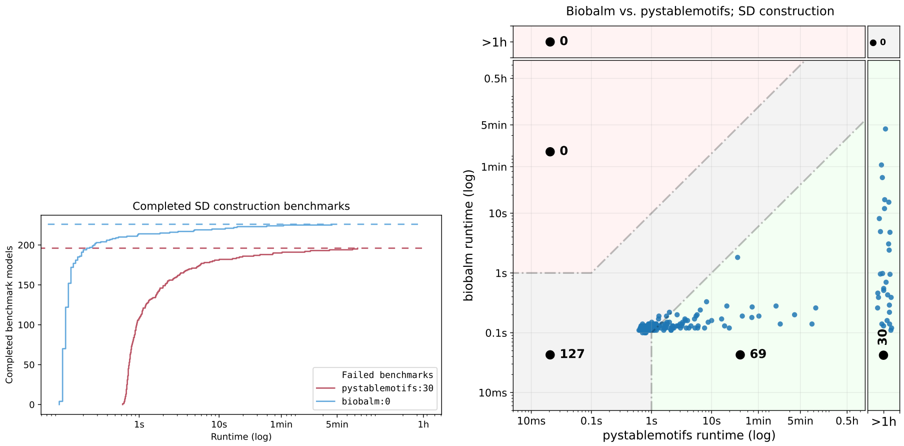


Figure S9: Performance comparison of succession diagram construction between **biobalm** and **pystablemotifs** on a the simplified BBM dataset with all input nodes fixed to 1.

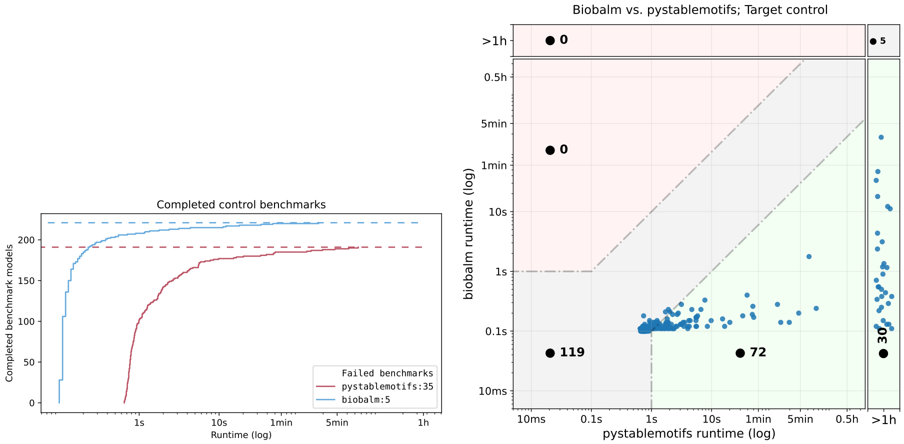


Figure S10: Performance comparison of computing control strategies for a single minimal trap space in **biobalm** and **pystablemotifs** on a the simplified BBM dataset with all input nodes fixed to 1.

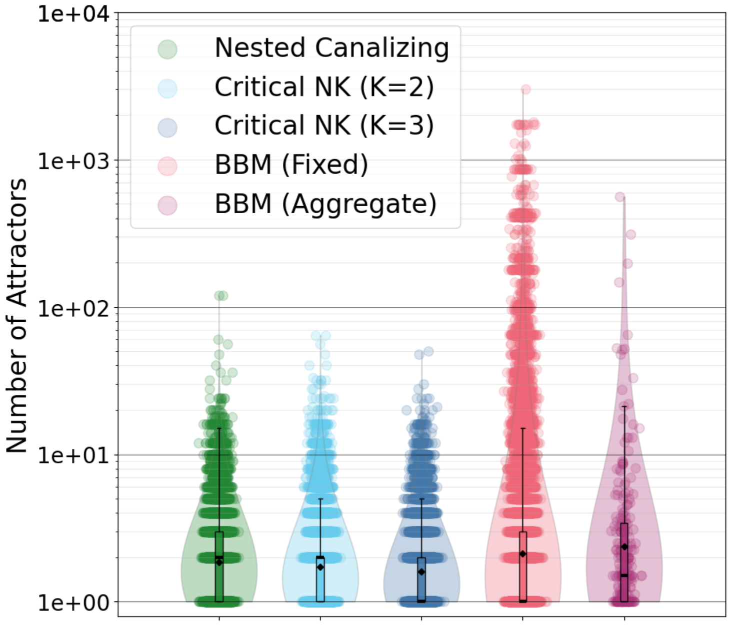


Figure S11: Distributions of number of attractors (also equal to the number of minimal trap spaces) for various ABN ensembles. BBM networks with randomly selected input configurations (up to 128 independent samples) are shown in red, with the average number of attractors across these samples for each network depicted in purple. Random networks of different types are indicated in green, cyan, and dark blue, and are constructed to match the distribution of the number of variables in the BBM ensemble. Gaussian noise is added to the horizontal position of each point. Box plots and density plots are computed in log space.

| number of nodes in the fully expanded succession diagram | | | | |
|---|---|---|---|---|
| | NCF | NK2 | NK3 | BBM_agg |
| mean | 2.01<br>(2.00, 2.03) | 1.70<br>(1.68, 1.71) | 1.76<br>(1.74, 1.78) | 2.41<br>(2.08, 2.80) |
| variance | 1.64<br>(1.61, 1.67) | 1.78<br>(1.75, 1.82) | 1.78<br>(1.74, 1.81) | 7.86<br>(5.59, 11.42) |
| kurtosis | 0.41<br>(0.28, 0.62) | 0.36<br>(0.23, 0.53) | 0.34<br>(0.21, 0.54) | 5.68<br>(3.46, 8.82) |

| number of attractors | | | | |
|---|---|---|---|---|
| | NCF | NK2 | NK3 | BBM_agg |
| mean | 0.90<br>(0.88, 0.91) | 0.78<br>(0.77, 0.79) | 0.67<br>(0.66, 0.68) | 1.25<br>(1.06, 1.49) |
| variance | 0.81<br>(0.79, 0.82) | 0.77<br>(0.75, 0.79) | 0.67<br>(0.65, 0.68) | 2.59<br>(1.88, 3.80) |
| kurtosis | 0.39<br>(0.19, 0.73) | 0.74<br>(0.56, 0.99) | 0.76<br>(0.59, 0.99) | 5.17<br>(2.98, 9.51) |

Table S2: Mean, variance, kurtosis of the number of nodes in the fully expanded succession diagram and the number of attractors, when logged with base 2. 95% confidence intervals of the statistics, obtained with 'BCa' method of the bootstrap method of the `scipy` library, is in parentheses. The confidence interval of the statistics of the BBM ensemble (averaged over the sampled configurations) does not overlap with the intervals of any of the random ensembles.

#### S7.3 Statistical test

We used bootstrap method to test the statistical significance of the difference between the succession diagrams in the random and the BBM ensemble. We tested the distribution of the number of nodes in the fully expanded succession diagrams and the number of attractors. In both distributions, the difference of the mean, variance, and kurtosis were statistically significant between the BBM ensemble and any one of the random ensemble. For each BBM model, we generated 100 models of the same size for each random ensemble, which results in 23000 models for each random ensemble. We constructed the fully expanded succession diagram for each of the networks, and found the number of nodes in the succession diagram, its depth, and the number of attractors. For the BBM networks, we randomly sampled up to 128 input configurations, and found the average value of the aforementioned numbers. We applied 'BCa', 'percentile' and 'base' method of the bootstrap method in `scipy` library on the logged values to find the 95 percent confidence interval of each statistial measure. The bootstrap distribution was plotted in each case to check the validity of the bootstrap method. In all 3 methods, the confidence intervals of each measure for BBM model ensemble did not overlap with any of the confidence intervals for the random ensembles. The result was similar when we did not take the log of the values.

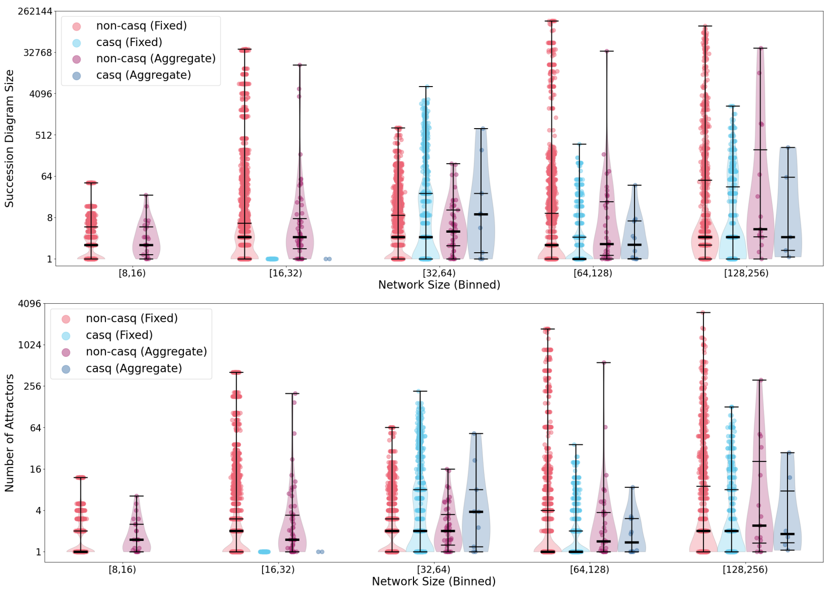


Figure S12: CaSQ and non-CaSQ models did not show significant difference in terms of the distribution of the number of nodes in the fully expanded succession diagram, or in terms of the distribution of the number of attractors. They shared the same long-tail behavior.

#### S7.4 CaSQ and non-CaSQ models

Here, we show that the long-tailed behavior of the BBM ensemble does not come from the fact that BBM also contains machine generated models. CaSQ [49] is a python package that automatically converts a molecular interaction graph to an executable Boolean model. In the BBM ensemble, we have 29 networks constructed with CaSQ. Figure S12 shows that the CaSQ and non-CaSQ models do not show significant difference in terms of the long-tail behavior in both the distribution of the number of nodes in the fully expanded succession diagram and in the distribution of the number of attractors.

#### S7.5 Depth scaling of SD size and attractors

Figure S13 shows that the number of attractors grows exponentially as the succession diagram depth increases. It demonstrates that the number of attractors is roughly bounded by $2^d$. Note that the succession diagram is free to have any depth, any size, and any number of attractors, and hence this rough upper bound comes from the property of the ensemble. It is noteworthy that these upper bounds follow the scaling of networks of independent cycles. Every independent cycle (e.g., a source cycle) increases the SD depth by 1 and doubles the number of attractors. Simple calculation shows that the SD size triples with

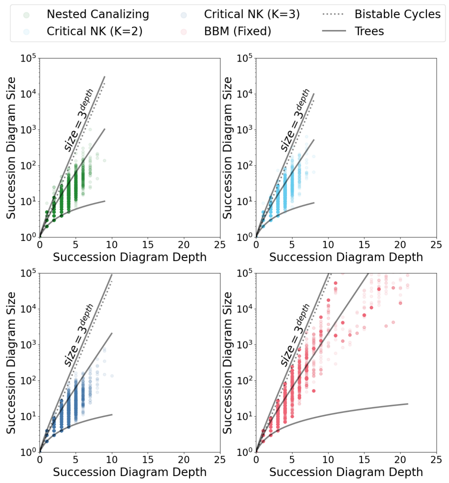


(a) Succession diagram size scales exponentially as succession diagram depth increases. Reference lines are given corresponding to the scaling for trees with 1, 2, or 3 children per node (solid). Additionally, the scaling for networks consisting of independent bistable cycles is shown with a dotted line.

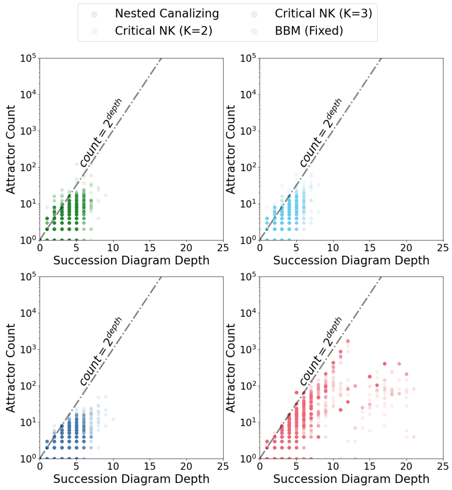


(b) Maximum attractor number scales approximately exponentially with succession diagram depth. The reference line correspond to succession diagrams that are trees with two children per node and also to (non-tree) succession diagrams generated by ABNs composed entirely of independent bistable cycles.

Figure S13: Succession diagram scaling within the BBM ensemble.

every added independent cycle.

### S8 Others

In this section, we list further data that supports the contents of the main paper, but does not require additional context or discussion.

#### S8.1 Toy example no. 2

Update rules of the second example network used in the main paper, together with its state-transition graph.

$$
\begin{aligned}
f_{v_1}(x) = \;& x_1 \vee (x_2 \wedge x_3) \\
f_{v_2}(x) = \;& x_2 \wedge \neg x_3 \\
f_{v_3}(x) = \;& (x_2 \wedge \neg x_3) \vee \\
& (\neg x_2 \wedge x_3 \wedge x_4) \vee \\
& (\neg x_2 \wedge \neg x_3 \wedge \neg x_4) \\
f_{v_4}(x) = \;& \neg x_1 \wedge [(x_3 \wedge x_4) \vee \\
& (\neg x_2 \wedge \neg x_3 \wedge \neg x_4)]
\end{aligned}
$$

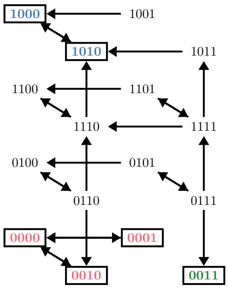


##### S8.2 Toy example no. 3

Update rules of the third example network used in the main paper, derived from model BBM-188 [50]. The derived model is constructed by setting all input nodes to *true*, percolating their values, and then reducing (inlining) network variables `Sox9_p_b2`, `Fgf9_p_b2`, `Fgf9_cr_b2`, `Fgf9_c_b2`, `Sox9_c_b2`, `Sox9_c_b1`, `Fgf9_cr_b1`, `Sf1_c_b2`, `Sry_c`, `Sry_p`, `Sf1_p_b2`. Variable indices are then assigned to the remaining variables alphabetically. We manually verified that this reduction does not change the succession diagram in any way.

$$
\begin{aligned}
f_{v_1} &= \neg v_2 \\
f_{v_2} &= \neg v_1 \\
f_{v_3} &= \neg v_2 \\
f_{v_4} &= (\neg v_3 \wedge v_6) \vee (v_3 \wedge v_2 \wedge v_6) \vee (v_3 \wedge \neg v_2) \\
f_{v_5} &= v_4 \wedge v_6 \wedge v_7 \wedge \neg v_8 \\
f_{v_6} &= v_4 \vee (v_6 \wedge v_5) \\
f_{v_7} &= v_7 \vee \neg v_8 \\
f_{v_9} &= \neg v_8 \\
f_{v_8} &= \neg v_9
\end{aligned}
$$

[26] James Lyle Peterson. *Petri net theory and the modeling of systems.* Prentice Hall PTR, 1981.

[27] David Angeli, Patrick De Leenheer, and Eduardo D. Sontag. Persistence results for chemical reaction networks with time-dependent kinetics and no global conservation laws. *SIAM J. Appl. Math.*, 71(1):128–146, 2011.

[28] Elisabeth Degrand, François Fages, and Sylvain Soliman. Graphical conditions for rate independence in chemical reaction networks. In *Proc. of CMSB*, pages 61–78. Springer, 2020.

[29] GaiYun Liu and Kamel Barkaoui. A survey of siphons in Petri nets. *Inf. Sci.*, 363:198–220, 2016.

[30] Olivier Coudert and Jean Christophe Madre. A new method to compute prime and essential prime implicants of Boolean functions. In *Advanced Research in VLSI and Parallel systems*, pages 113–128. The MIT Press, 1992.

[31] Van-Giang Trinh, Belaid Benhamou, Thomas Henzinger, and Samuel Pastva. Trap spaces of multi-valued networks: definition, computation, and applications. *Bioinf.*, 39(Supplement_1):i513–i522, 2023.

[39] Samuel Pastva, David Safranek, Nikola Benes, Lubos Brim, and Thomas Henzinger. Repository of logically consistent real-world Boolean network models. *bioRxiv*, 2023.
